# Dynamic adaptation of sequential action benefits from cortico-basal ganglia-related temporal variability

**DOI:** 10.1101/2022.03.28.486040

**Authors:** Lachlan A. Ferguson, Miriam Matamales, Bernard W. Balleine, Jesus Bertran-Gonzalez

## Abstract

Performing several actions in swift succession is often necessary to exploit known contingencies in the environment. However, in order to remain successful when contingency rules change, streamlined action sequences must be adaptable. Here, by combining analyses of behavioural microstructure with circuit-specific manipulation in mice, we report on a relationship between action timing variability and successful adaptation that relies on post-synaptic targets of primary motor cortical (M1) projections to dorsolateral striatum (DLS). Using a two-lever instrumental task, we found that mice build successful action sequences by first establishing action scaffolds, from which they dynamically elongate as task requirements extend. Specific interruption of the M1→DLS circuit altered these dynamics, prompting actions that were less variable in their timing, overall reducing opportunities for success. Our results reveal a role for M1→DLS circuitry in setting the exploration/exploitation balance that is required for adaptively guiding the timing and success of instrumental action. Based on evidence from transsynaptic tracing experiments, we propose that such function may involve additional downstream subcortical processing relating to collateralisation of descending motor pathways to multiple basal ganglia centres.

## Introduction

For stable environmental contingencies that require more than a single action, animals can learn to perform a series of discrete responses that, as experience accrues, melds the internal boundaries between actions into accurately timed streams of skilled behaviour, often expressed as a single unit or “chunk” (Graybiel, 1998; Lashley, 1951; Rosenbaum et al., 1983; Saling & Phillips, 2007; Sternberg et al., 1978; Terrace, 1987). In a fluctuating environment, however, behavioural streams must incorporate a sustained degree of variability if they are to remain adaptive (Sternad, 2018). Animals can show a dramatic variability in the rate of responding that is specifically promoted by the external requirements of the task—such as reinforcement schedule and session duration—rather than by internal motivational states (Dezfouli et al., 2019; McSweeney & Roll, 1993). It has been argued that this variability in performance (or within-organism “noise”) can sustain itself through reinforcement (known as ‘reinforced variability’), such that fluctuating behaviour is instrumental in achieving outcomes (Neuringer, 2002). Importantly, behavioural noise is multifaceted in at least 2 ways: (i) the variation of the number of actions, and (ii) the variation in the timing of these actions—both of which have been shown to contribute to establishing optimal performance (Light et al., 2019). Such forms of behavioural variability therefore constitute a valuable source of change that can support the adaptation of well-established sequential action when environmental conditions demand it.

The modification of well-established action appears to be particularly dependent on motor cortical inputs targeting underlying basal ganglia structures. For example, during the development of overtrained actions, such as habits and skills, the functional priority of cortical projections to distinct basal ganglia regions appears to be reorganised from medial prefrontal cortex and dorsomedial striatum (DMS) to sensorimotor cortices and dorsolateral striatum (DLS) (Balleine, 2019; Balleine et al., 2007; Balleine & O’Doherty, 2010; Ostlund & Balleine, 2005; Yin et al., 2004, 2006). In the context of motor skill learning, corticostriatal afferents targeting the DMS and DLS initially co-engage, but as skills develop, medial associative input strength declines more rapidly and to a greater degree than lateral motor cortical inputs (Kupferschmidt et al., 2017). Beyond major intratelencephalic corticostriatal projections, several recent studies have highlighted the functional importance of motor corticofugal systems collateralising over underlying basal ganglia nuclei, such as the DLS, the globus pallidus externa (GPe) and the subthalamic nucleus (STN) (Karube et al., 2019; Kita & Kita, 2012; Nelson et al., 2021), altogether opening the door to additional sources of bottom-up motor output processing.

In the striatum, studies recording neuronal activity or assessing function using chemogenetics implicate lateralised striatal regions in the regulation of action chunking and timing (D. Z. Jin et al., 2009; X. Jin et al., 2014; X. Jin & Costa, 2010; Matamales et al., 2017; Mello et al., 2015). Contributions to the temporal dynamics of well-learned action sequences have also been observed following manipulations of direct pathway projection neuron activity in the DLS (e.g., optogenetic stimulation extends ongoing action sequences) (Tecuapetla et al., 2016), whereas chemogenetic inhibition during learning compresses sequences into briefer durations without impacting the total number of presses within them (Matamales et al., 2017). Similarly, pharmacological DLS inactivation in well-trained animals can preserve action sequence structure while increasing trial-by-trial variability (Rueda-Orozco & Robbe, 2015). Collectively, while evidence strongly supports the contribution of cortical projections directly or secondarily targeting DLS neurons in establishing both the structure and timing of well-learned sequences, the way the downstream circuitry promotes further adaptation of action remains unstudied.

Here, we hypothesised that DLS postsynaptic processing of motor cortical information shapes action sequence structure by accommodating temporal variability, a process that is critical for successfully adapting ongoing sequential action. By combining circuit-specific cell ablation with an instrumental paradigm that specifically promotes self-determined variation of action timing, we identified that mice perform sequences with stable inter-press intervals following initial sequence acquisition, then smoothly integrate further behavioural segments into larger action sequences when contingency requirements increase. We found that a subtle depletion of M1→DLS post-synaptic connectivity destabilises this process by reducing timing variability, something that, we propose, could rely on motor corticofugal circuitry collateralising to the basal ganglia.

## Results

### As mice build skilled actions, the timing of their sequences adapts with success

The first experiment aimed to characterise the behavioural adaptations that mice make to the timing and efficacy of their performance over instrumental training. To ensure that action timing and performance remained self-paced and uninfluenced by external cues common in instrumental conditioning procedures (e.g., pellet delivery sounds), we developed a self-paced chained-sequence task based on the Mechner Counting Task (Mechner, 1958; Light et al., 2019). Mice were presented with two levers in tandem and earned a reward for a single press on the second ‘End’ lever, provided they had completed the required number of presses on the first ‘Sequence’ lever (Figure 1A, top). This allowed the mice to freely decide the duration of their action chains (performed on the Sequence lever) without relying on external cues associated with reward delivery. The requirements on the Sequence lever increased every four sessions from fixed ratio (FR) 1 to FR3 to FR5 across training, whereas the End lever always required one single press (Figure 1A, bottom). General measures of performance in this task, such as lever press rate (Figure 1B) and number of presses per sequence (Figure 1C), significantly increased for the Sequence lever but not for the End lever over the course of training after pretraining (Figure 1–figure supplement 1A). This was supported by a significant session x lever interaction in both cases (Table supplement 1), demonstrating that mice appropriately biased performance toward the Sequence lever. Mice also significantly increased the rate of rewards earned within each FR schedule (Figure 1D and Table supplement 1), which was accompanied by a commensurate reduction in the number of sequences required to achieve them (Figure 1E), as well as reduced magazine entry rates (Figure 1–figure supplement 1B and Table supplement 1). These results show that mice clearly distinguished between lever contingencies and adapted well in each phase of the task.

**Figure 1.**
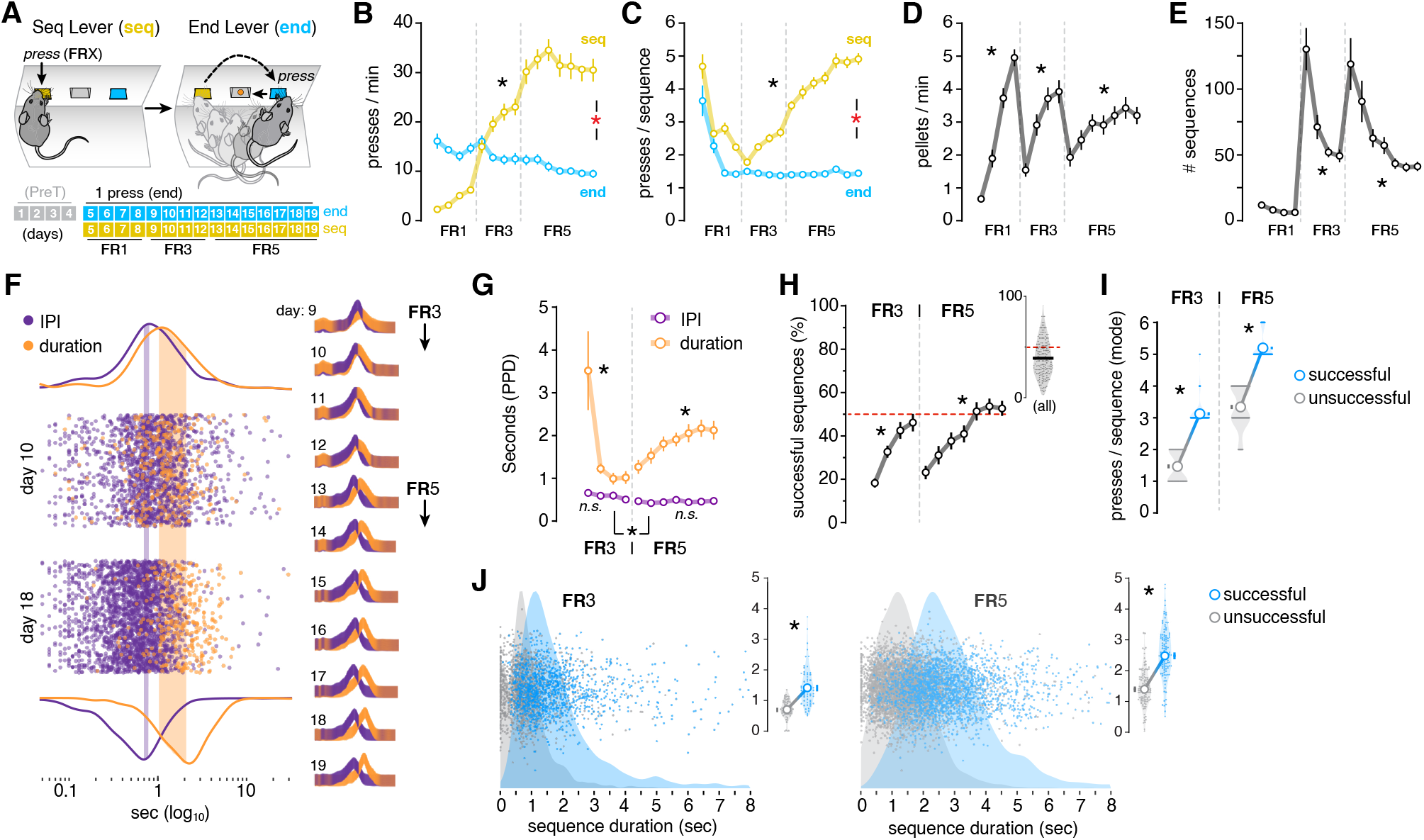
A self-paced sequence task reveals timing adaptations during training. (**A**) Animals were pre-trained with continuous reinforcement (1 press→1 reward) on the End lever for 4 sessions (sessions 1-4, PreT, see Figure 1–figure supplement 1A). Next, the Sequence lever was introduced on a fixed ratio (FR) 1 schedule for 4 sessions (sessions 5-8), whereby pressing on the Sequence lever must occur prior to pressing the End lever in order to receive reward. In the following 4 sessions (sessions 9-12), the press requirements on the Sequence lever increased to FR3. In the final 7 sessions (sessions 13-19), the press requirements on the Sequence lever increased to FR5. (**B**) Lever press rate measured as presses per minute on each lever type throughout FR1-FR5 training. (**C**) Sequence length measured as the number of presses per sequence in both the Sequence and End levers across FR1-FR3 training. (**D**) Reward rate measured in pellets per minute. See Figure 1–figure supplement 1C for the total rewards earned throughout training. (**E**) Total number of sequences performed throughout FR1-FR5 training. (**F**) Scatter plot of each IPI and sequence duration value (to log_10_) on the Sequence lever for all animals in an example FR3 (day 10) and FR5 (day 18) session, with probability density function curves indicating peak differences (shaded). Right diagrams show the probability density function curves on each day of FR3-FR5 training. See Figure 1–figure supplement 2A-B for individual days. (**G**) IPI and Duration expressed as averaged probability density peaks (PPD, seconds) across FR3 and FR5 training. (**H**) Percentage of sequences that successfully resulted in reward in FR3 and FR5 sessions across training and in all FR3 and FR5 training sessions collapsed (inset). Red dashed line denotes 50%. (**I**) Most frequently occurring (modal) number of presses in either unsuccessful or successful sequences for both FR3 or FR5 training. Truncated violin plots are fitted to data points (shaded). (**J**) Scatter plot with probability density function curves (shaded) of sequence duration for every unsuccessful (unrewarded) and successful (rewarded) sequence performed by the entire cohort during FR3 (left) and FR5 (right). Insets show PPD for each animal and day during FR3 (left) and FR5 (right) training. *, significant overall/simple effect (black) and interaction (red). N.S., not significant (Table supplement 1).

We then assessed if the improved effectiveness in earning rewards coincided with more efficient action sequence timing. To determine what type of changes in timing predominated during the adaptation of action, we measured the peak probability of both individual inter-press intervals (IPIs) within a sequence and whole sequence durations across training (FR3 and FR5). We found that the probability density distribution for IPIs remained stable throughout training, whereas the same function applied to sequence duration shifted to the right as training progressed (Figure 1F and Figure 1–figure supplement 2A and B). Measures of the action timing peak distribution across training revealed that IPIs remained relatively stable, with only a moderate decline occurring between FR3 and FR5 phases. In contrast, the sequence duration initially declined during early acquisition (FR3), then steadily increased during FR5 training (Figure 1G and Table supplement 1). Given the rise in reward rate, the related decline in the number of sequences, and the elongation of the number of presses per sequence, we expected the likelihood of performing action sequences that ended in reward to increase as training progressed. We calculated the percentage of successful (rewarded) sequences relative to unsuccessful (unrewarded) sequences and found that the former significantly increased within each training phase (FR3 and FR5), reaching 39.11% on average across all training and, after five sessions, plateauing at approximately 50% success on FR5 (Figure 1H and Table supplement 1). Mice were clearly capable of improving the efficacy of their sequences by increasing the chance of performing—at minimum—the required number of presses. It was unclear, however, if the adjustments to the number of actions from unsuccessful to successful trials coincided with adjustments in sequence timing; i.e., whether (i) more presses were added to a fixed period and executed at a faster rate, or (ii) sequence duration was extended with the addition of lever presses executed at a similar rate. We observed that the latter was the case: when a significantly greater number of presses was implemented for successful sequences (Figure 1I and Figure 1–figure supplement 1D), the peak sequence duration of successful sequences shifted to significantly longer durations relative to unsuccessful attempts (Figure 1J and Figure 1–figure supplement 1E, Table supplement 1). These data suggest that when it comes to action timing, over and above changes in inter-press-intervals, the modulation of sequence duration seems to be the critical variable when adapting action for success.

### New chunks are smoothly merged with previously established action scaffolds

We next examined how successful sequences are constructed when facing a change in schedule. We investigated this by quantifying the frequency of all sequences according to the number of presses per sequence (Figure 2A and B). Then, to discern the likelihood of performing a sequence comprised of a given number of presses for either sequence category (i.e., unsuccessful and successful), we independently calculated the probability for each total number of presses per sequence that were performed in either successful or unsuccessful sequences (Figure 2A and B, insets). Interestingly, we found that successful responses were often a 2-press chunk away from the most frequent unsuccessful sequence; i.e., responses frequently transitioned from unsuccessful 1-press actions to successful 3-press sequences in FR3 training (Figure 2A), and from unsuccessful 3-press sequences to successful 5-press sequences in FR5 training (Figure 2B). This analysis showed that, on experiencing increases in contingency, mice demonstrated an ability rapidly to modify their sequences to match the new schedule requirements without excessive underand overshooting.

**Figure 2.**
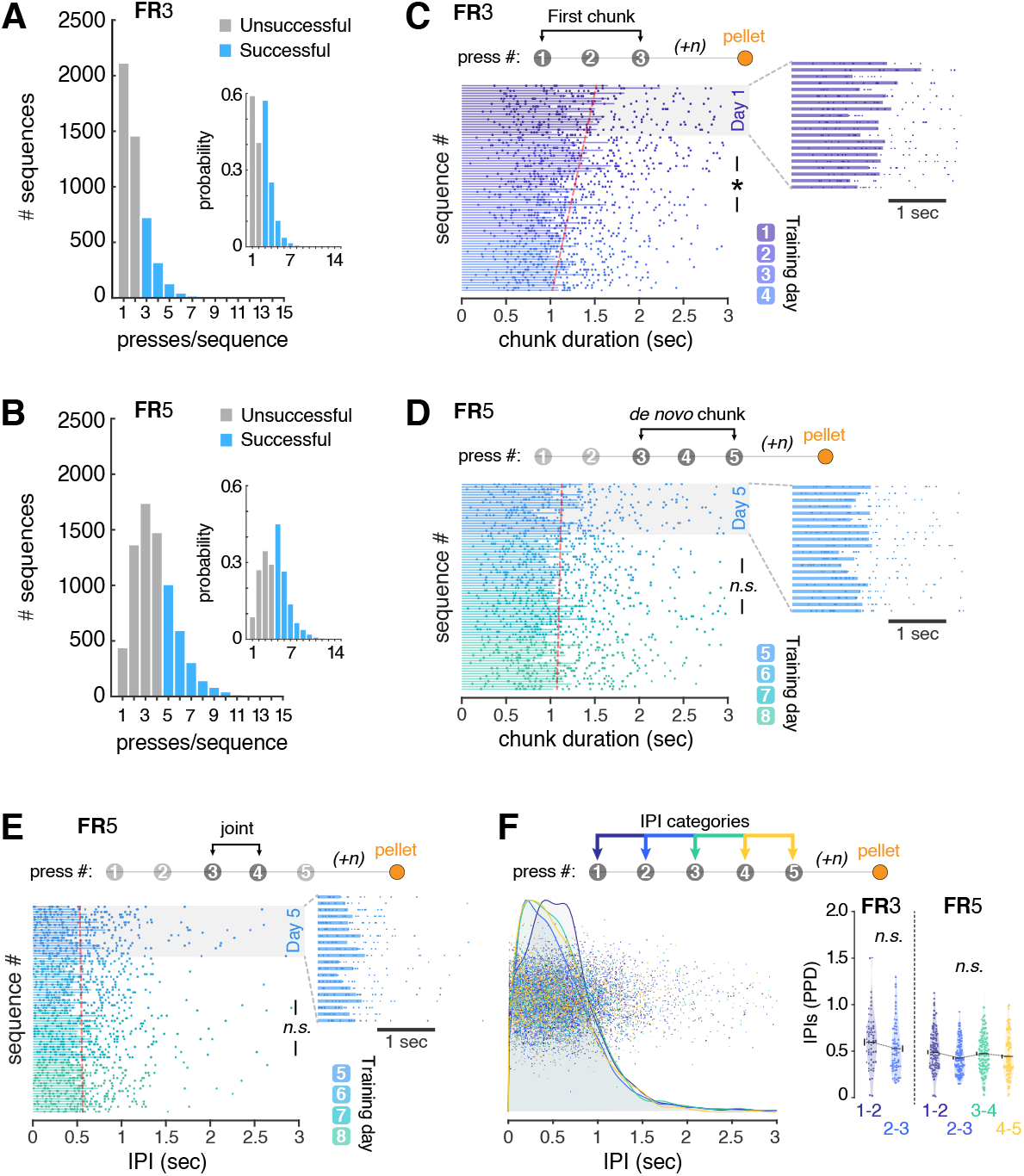
Successful action dynamics smoothly evolve as task requirements increase. (**A-B**) Frequency histograms showing the total number of sequences performed on the sequence lever according to the number of presses per sequence during FR3 (A) and FR5 (B) training. Insets show the probability distribution of the same sequence categories during FR3 (A, right) and FR5 (B, right) training. (**C-E**) Duration of successful subsequence intervals ranging from presses 1-3 (First chunk, C), presses 3-5 (*de novo* chunk, D) and presses 3-4 (joint, E) arranged chronologically across the first four sessions of FR3 and FR5. Data are the duration of each sequence by each mouse (dots) and the average across mice (bars). A linear regression model highlighting the chronological trend is fitted to the data (red dashed line). Insets are an enlarged view of the first session of the corresponding fixed ratio schedule. (**F**) Scatter plot with probability density function curves (shaded) of IPIs between 1-2, 2-3, 3-4 and 4-5 press transitions for every successful sequence performed by the entire mouse cohort during FR3 and FR5 training (left). Peak probability density (PPD) of relevant IPIs in successful sequences for each training session (dots) plotted for both FR3 and FR5 (right). n = any number of presses before reward. *, significant overall/simple effect (black) and interaction (red). N.S., not significant (Table supplement 1).

Next, we explored the way lever press responses were chunked during action sequence learning and whether temporal gaps between chunks emerged as animals adapted their sequences to new ratio requirements (Rosenbaum et al., 1983). For this, the duration of each chunk within successful sequences was arranged chronologically (following the order in which each sequence occurred within a session). To observe the relationship between the chronological progression of successful sequences and the duration of their constituent chunks, we analysed their linear relationship over both FR3 and FR5 training (Figure 2C and D, Figure 2–figure supplement 1). We found that during FR3 training—when sequences are first acquired—the duration of the first chunk (time between presses 1-3) significantly declined over training (Figure 2C and Figure 2–figure supplement 1A, Table supplement 1). In contrast, during FR5 training—when FR3 sequences have already been established and two extra presses are being added—the duration of the *de novo* chunk (time between presses 3-5) was the same as the first chunk in late FR3 training (∼1 sec; Figure 2D), and remained constant throughout the rest of training (Figure 2D and Figure 2–figure supplement 1B, Table supplement 1). In light of the observed disparities in the evolution of the first and *de novo* chunks of successful sequences, we investigated if the two chunks were implemented as discrete units with a pause between them, or if they were smoothly integrated into an extended single sequence of action. We found that the time in-between the two chunks (i.e., the “joint” IPI; between presses 3 and 4) remained invariable as rewarded experience accrued across FR5 training (Figure 2E and Figure 2–figure supplement 1C, Table supplement 1). Furthermore, we found that the different IPI categories across successful sequences were mostly indistinguishable from each other, including the joint IPIs connecting first and *de novo* chunks (Figure 2F, Table supplement 1). Collectively, these data reveal that mice smoothly integrate new sub-sequence chunks with previously acquired sequence prototypes to immediately form extended sequences.

### Circuit-specific interruption of the M1→DLS corticostriatal pathway speeds up sequential action

Functional assays, such as lesion and chemogenetic suppression, indicate that the DLS governs a variety of roles relevant to optimising task performance in sequence-based instrumental conditioning, ranging from skilled action kinematics, speed and variation of action sequences, habit learning and the accurate acquisition of a serial order (Dhawale et al., 2021; Jurado-Parras et al., 2020; Matamales et al., 2017; Rueda-Orozco & Robbe, 2015; Yin, 2010; Yin et al., 2004). Similarly, the M1 and its connectivity with the DLS has also been implicated in the acquisition and governance of the constituent components of skilled action sequences (Kawai et al., 2015; Kupferschmidt et al., 2017; Nelson et al., 2021). We investigated whether M1 projections in specific post-synaptic targets of the striatum contributed to modulating action time during sequence learning, and whether this influenced task success. In adult mice, we interrupted the M1→DLS pathway through an AAV-based circuit-specific lesion approach, which combined anterograde transport of Cre with Cre-dependent lesion (Gradinaru et al., 2010; Yang et al., 2013) (Figure 3A). A first AAV expressing anterograde travelling, trans-synaptic WGA-Cre (AAV2-EF1a-mCherry-IRES-WGA-Cre; Antero-Cre) was injected into the M1

**Figure 3.**
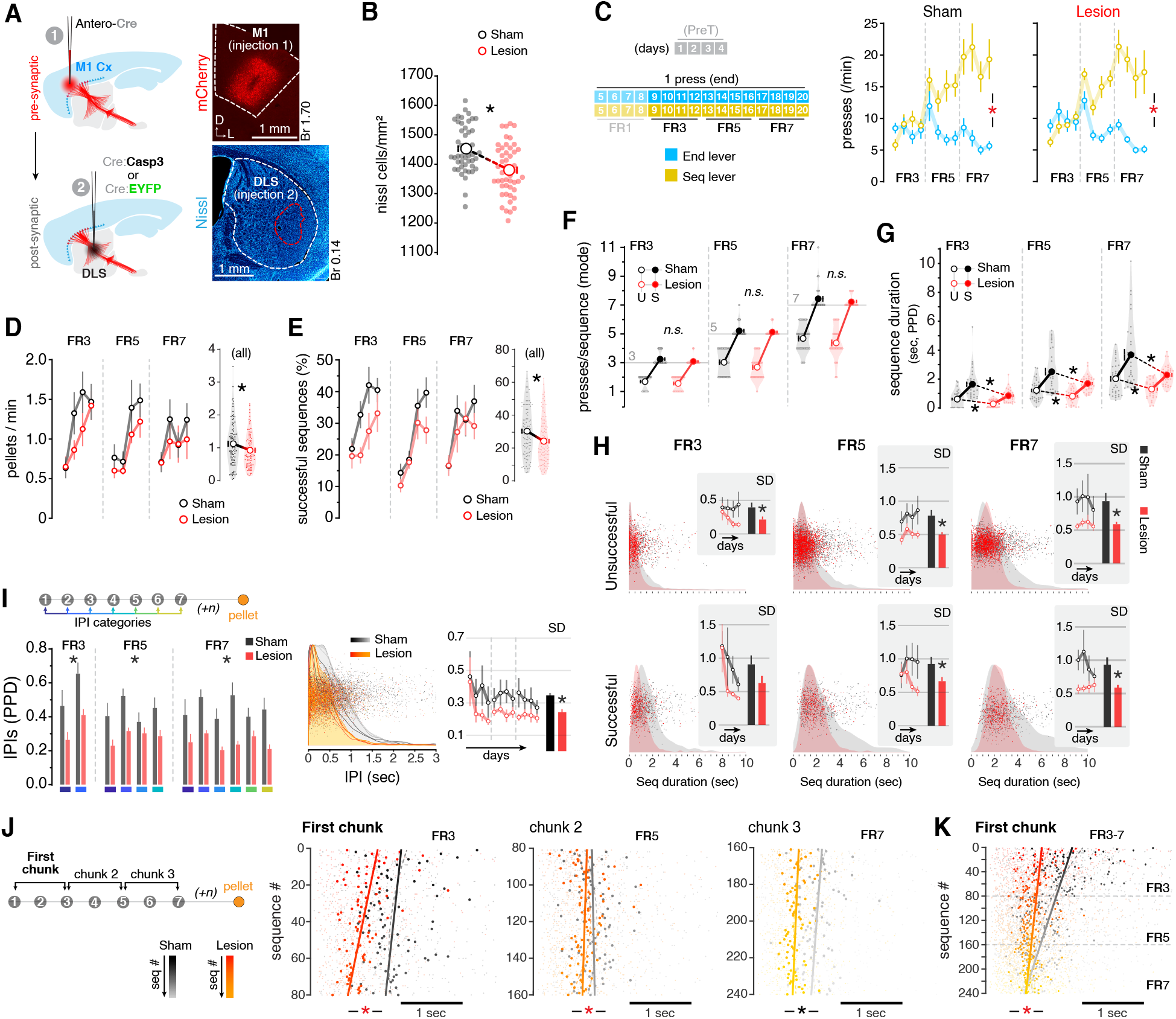
Specific interruption of the M1→DLS circuit alters the temporal dynamics of action sequences. (**A**) Schematic of the M1→DLS circuit lesion strategy. (1) An anterograde AAV expressing Cre (Antero-Cre) was injected into the M1. (2) An AAV expressing either Cre-dependent synthetic procaspase taCasp3 (Cre:Casp3; Lesion group) or Cre-dependent EYFP (Cre:EYFP; Sham group) was then injected into the DLS. Top-right: confocal micrograph showing expression of Antero-Cre virus in the M1 injection site revealed by mCherry. Bottom-right: confocal micrograph showing the usual extent of the lesion at the Cre:Casp3 injection site in the DLS revealed by Nissl labelling. (**B**) Nissl-based cell density quantification within DLS injection site. (**C**) Top: animals began pre-training with continuous reinforcement (CRF) on the End lever (Lend) for 4 sessions (PreT), then shifted to the tandem task on the Sequence lever (Lseq) progressing through FR1→FR3→FR5→FR7 schedules every 4 sessions (top). Bottom: lever press rate measured as presses per minute on each lever type throughout FR3→FR7 training in each group. For sessions 1-4 (PreT), see Figure 3–figure supplement 4C. (**D**) Reward rate (press/min) throughout FR3→FR7 training. Inset shows data from the 3 schedules collapsed. (**E**) Percentage of successful sequences across FR3→FR7 training. Inset shows data from the 3 schedules collapsed. (**F**) Most frequent (modal) number of presses in a sequence (press/sequence) for both Successful and Unsuccessful sequences in FR3, FR5 and FR7 schedules. (**G**) Sequence duration (peak probability distribution, PPD) for both Successful and Unsuccessful sequences in FR3, FR5 and FR7 schedules. (**H**) Scatter plot with PPD curves (shaded) of sequence duration for every unsuccessful (top) and successful (bottom) sequence produced by the entire cohort during FR3, FR5 and FR7 phases. Insets show standard deviation (SD) across training sessions and a summary bar graph of all sessions within the indicated schedule. (**I**) Left: schematic for quantification of IPIs within a sequence. Data (bottom) shows the PPD of the relevant IPIs for successful sequences in each schedule. Centre: Scatter plot with PPD functions (shaded) of IPIs between the indicated press transitions for every successful sequence performed by the entire cohort as FR3→FR7 training progressed (colour coded). Right: standard deviation (SD) across FR3→FR7 training. Inset shows a summary bar graph of all sessions collapsed. (**J**) Duration of successful subsequence intervals ranging from presses 1-3 (First chunk, left), presses 3-5 (chunk 2, centre) and presses 5-7 (chunk 3, right) arranged chronologically across FR3, FR5 and FR7 sessions. Data are the duration of each sequence by each mouse (small dots) and the average across mice (larger dots). A linear regression model highlighting the chronological trend is fitted to the data (line). The change in colour in each group’s dataset reflects progression throughout training (see legend to the bottom-left). (**K**) Duration of the first successful subsequence segment (first chunk) throughout the entire FR3→FR7 training. n = any number of presses before reward. *, significant overall/simple effect (black) and interaction (red). N.S., not significant (Table supplement 1).

(Figure 3–figure supplement 1A), followed by delivery of a second AAV into the DLS, which expressed either Cre-dependent procaspase 3 (AAVFlex-taCasp3-TEVp; Cre:Casp3) or Cre-dependent EYFP (AAV5-EF1A-DIO-eYFP; Cre:EYFP) (Figure 3–figure supplement 1B). Cre-Casp3-injected mice (Lesion group) showed a significant reduction of neuronal density in a defined area of the DLS compared to Cre:EYFP mice (Sham group) (Figure 3B). When exposing these mice to the self-paced sequence task, we found that mice from both the Lesion and Sham groups appropriately biased lever press performance toward the Sequence lever as sequence training progressed from FR3 to FR7 (Figure 3C), revealed by a strong session x lever interaction in both groups (Table supplement 1). On the other hand, a summary of task success across all sequence training (FR3-FR7) showed that M1→DLS lesioned mice earned rewards at a slower rate (Figure 3D, Table supplement 1) and performed sequences with a lower percentage of success relative to Sham controls (Figure 3E, Table supplement 1), without impacting the total number of earned rewards or magazine entries per session (Figure 3–figure supplement 1D and E). Despite this reduced percentage of successful actions, the M1→DLS lesioned group showed no difference in the number of presses per sequence relative to the Sham group when performing either unsuccessful or successful sequences (Figure 3F, Table supplement 1). By contrast, the timing of these sequences was altered, such that both unsuccessful and successful sequences were faster in M1→DLS lesioned mice (Figure 3G, Table supplement 1).

### The M1→DLS pathway supplies action timing variability required for adaptation

Next, we explored the relationship between sequence speed increases and the variability of their execution as a source of explanation for the reduced success of M1→DLS lesioned animals. We found that the M1→DLS lesioned group maintained more consistent durations across unsuccessful and successful sequences compared to Sham controls throughout training (Figure 3H), the latter group showing significantly more variable sequence durations of either type at each phase of training (Table supplement 1). Consistent with a variability-based explanation of task success, within-subjects analysis showed that successful sequences were significantly more variable than unsuccessful sequences in both groups (Figure 3–figure supplement 1F, Table supplement 1). Further linear regression analysis showed that while successful sequence duration variability declined as task success increased for both groups, the slope of such decline was significantly less pronounced in M1→DLS lesioned mice (Figure 3–figure supplement 1G, Table supplement 1).

We then sought to clarify if, in successful sequences, the limited timing repertoires promoted by M1→DLS lesions generalised to whole sequence spans or if action timing limits were present in specific behavioural units within the sequence. By sorting IPIs according to position in the sequence and comparing their differences within each training schedule, we found a general significant decrease in IPI time following M1→DLS lesions across training, with no differences between the IPIs according to position in the sequence for either FR5 or FR7 training (Figure 3I, Table supplement 1). Moreover, similar to the variation reductions identified in whole sequences, a significant reduction in the variability of the IPIs in successful sequences was observed following lesion (Figure 3I-right panel, Table supplement 1). Further sequence structure analysis showed a reduced chunk duration for the first chunk (press 1-3) during FR3, for chunk 2 (press 3-5) during FR5 and for chunk 3 (press 5-7) during FR7 in the DLS lesion group, which significantly diverged from the Sham group as rewarded experience progressed in FR3 and FR5 (Figure 3J). Importantly, our analysis of the evolution of the first chunk—which we observed undergoes temporal change during initial action sequence acquisition in our previous experiment—revealed a highly supressed rate of change in M1→DLS lesioned mice, such that early training action speeds more closely resembled later training speeds relative to the significantly increasing speeds found in Sham controls (Figure 3K, Table supplement 1). Notably, this effect was not observed during the later acquisition of chunk 2 (press 3-5), or the duration of a successful sequence as a whole (Figure 3–figure supplement 1H and I). Overall, our results showed that M1→DLS interruption interfered with the successful construction and adaption of action sequences in response to an increasing lever press requirement whilst reducing the optimal range of action speed and variation.

### Motor cortical projections form multi-stage connections with the basal ganglia

An important consideration for disentangling M1→DLS function in adapting action sequence dynamics during learning is the likely involvement of subcortical bottom-up processing, something that, based on recent literature, could be promoted by collateralised connectivity in motor cortical descending pathways. For example, pyramidal tract neurons originating from layer V in the motor cortex are known to strongly collateralise to the subthalamic nucleus (STN), forming a “shortcut” into the basal ganglia commonly known as the hyperdirect pathway (Giuffrida et al., 1985; Nambu et al., 2002). Additionally, corticofugal projections are also known to emit accessory collaterals to more upstream basal ganglia structures such as the GPe or the striatum itself (Karube et al., 2019; Kita & Kita, 2012; Nelson et al., 2021). These collateral links to downstream basal ganglia nodes are thought to supply efferent copies of ongoing pyramidal tract motor commands, a process that can be key to adjusting the temporal limits of sequential action (Nambu et al., 2002; Nelson et al., 2021). We sought to clarify whether the motor cortical projections targeting the DLS pointed out in our study were also represented in downstream collateral networks of descending corticofugal pathways. We first explored the relative densities of motor cortical axons arborising through diverse basal ganglia structures using the Allen Mouse Brain Connectivity Atlas (Oh et al., 2014), which combines eGFP anterograde viral tracing with serial two-photon tomography throughout the entire brain. We selected 3 different cortical injection assays (spanning the M1, M2 and dorsal agranular insular area [AId] regions) based on their preference of projection to the STN (Figure 4A-C). EGFP-labelled axons in all three assays densely innervated several subcortical structures, particularly the DLS and lateral areas of the globus pallidus externa (lateral GPe) (Figure 4B). Quantification of EGFP fluorescence identified that, across major brain structures, the STN, striatum and GPe consistently had the three highest non-cortical projection densities over the 3 assays (Figure 4C).

**Figure 4.**
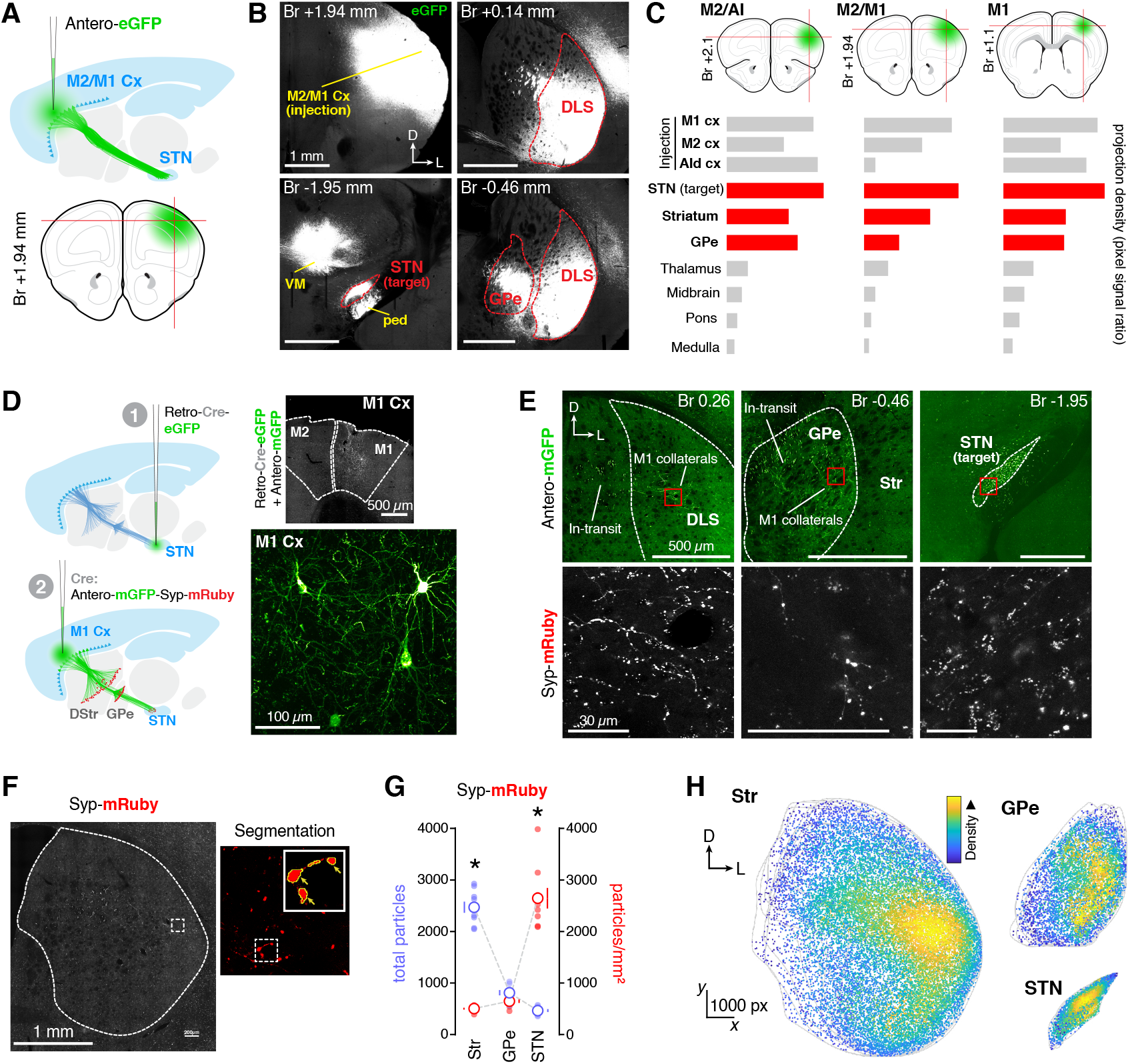
Motor corticofugal projections send shared collaterals to the DLS, GPe and STN. (**A-C**) Anterograde tracing study using the Allen Mouse Brain Connectivity Atlas database (Oh et al., 2014). Three different cortical injection assays targeting the STN were identified (Source: M1 and M2; Target: STN; see methods). (**A**) Experimental diagram on one of the assays showing the injection site onto M2/M1 cortex and expected transport of an anterograde reporter virus reaching the STN (antero-eGFP). (**B**) Example two-photon tomography images of eGFP expression in the M2/M1 injection site (top-left); the DLS (top-right); the STN (bottom-left) and the GPe (bottom-right) (experiment number 180709942). (**C**) Schematic for M2/agranular insular cortex (AI) (top-left), M2/M1 (top-centre), and M1 (top-right) injection sites with corresponding projection density quantifications throughout various brain areas. Data are extracted from experiments 180719293 (left), 180709942 (centre) and 100141780 (right). Projection densities for cortical regions around the injection site (‘Injection’) are listed first, followed by the three highest density regions (red)—including the site of target search (STN)—followed by other representative high-density regions. (**D**) Schematic depicting the viral tracing strategy used to identify hyperdirect pathway accessory targets: (1) a retrograde AAV expressing Cre-eGFP was injected in the STN. (2) an anterograde AAV expressing Cre-dependent mGFP and synaptophysin (Syp)-mRuby (labelling presynaptic boutons) was injected in the M1. Right panels are confocal images showing GFP expression in the M1. See Figure 4–figure supplement 1 for quantification of STN targeting. (**E**) Spinning disk confocal images of anterograde-mGFP in the striatum (top-left), GPe (top-centre) and STN (top-right); and Syp-mRuby-labelled terminals in each region (bottom panels). (**F**) Spinning disk confocal image showing Syp-mRuby clusters segmented for particle analysis (see methods). (**G**) Total particles and Particle density (particles/mm2) quantification for DStr, GPe and STN regions [*p<0.05]. (**H**) Particle density maps overlaid for each region (3x slice/animal; n = 3) on side ipsilateral to STN and M1 injection sites. See Figure 4–figure supplement 2 for individual maps.

We then investigated whether these regions (STN, striatum and GPe) were simply parallel cortical targets or in fact shared collaterals of the same corticofugal neurons. For this, we implemented a quantitative connectivity approach based on the retrograde transport of a Cre-expressing virus (AAV-hSyn-HI-eGFP-Cre-WPRE-SV40) injected at the most downstream target (STN, Figure 4–figure supplement 1), followed by anterograde transport of a Cre-dependent reporter virus (AAV-hSyn1-FLEX-mGFP-2A-synaptophysin-mRuby) injected at the origin of the corticofugal pathway (M1) (Figure 4D). Because the virus causes Cre-dependent anterograde expression of mRuby in synaptic terminals (Fisher et al., 2020; Zhang et al., 2016), this method allowed us to quantify the distribution of *any* mRuby-labelled synaptic boutons in projections collateral to and within the mainstream projection targeting of the STN. Consistent with the previous experiment, we detected dense mRuby puncta in both DLS and lateral GPe, in addition to the final target STN (Figure 4E). Our particle density analysis (Figure 4F) showed that the total number of mRuby+ synaptic boutons was greater in the collateral projections to the striatum and GPe compared to the STN, with the greatest number occurring in the striatum (Figure 4G, purple trace) (Table supplement 1). In contrast, the relative density of synaptic boutons within their regional space was greatest in the STN compared to the striatum and GPe (Figure 4G, red trace) (Table supplement 1). The reconstruction of the distribution of mRuby+ terminals into particle density maps showed that the greatest synaptic densities occurred within the lateral segments of the striatum and GPe, whereas synaptic territories remained central in the final target STN (Figure 4H, Figure 4–figure supplement 2). These data extend recent reports showing upstream cortico-basal ganglia connectivity (Karube et al., 2019; Kita & Kita, 2012; Nelson et al., 2021) and reveal that the cortical descending pathway originating in M1 indeed sends shared projections to the DLS. These results add to the growing literature emphasising accessory collateral networks of motor corticofugal systems as important players for the modulation of action timing.

## Discussion

### Leveraged adaptation of mature action sequences

A useful behavioural strategy for efficiently exploiting environmental contingencies often involves enacting accurately learned streams of swiftly executed actions. In fluctuating environments, however, sufficient variation in these streams of action must be generated for exploring newly adaptive solutions, a process described as ‘adaptive learning through variation and selection’ (Burtsev, 2012) in which, contrary to the decrease in variability typically observed with skill mastery, increased behavioural noise promotes adaptive learning (Sternad, 2018). Our study found evidence, amongst the varied successful responses, for a balance between effective exploration and efficient exploitation of instrumental action following an upward shift in contingency rules. As training progressed, the proportion of successful sequences within an action stream improved, and the probability of producing the target sequence (precisely at the required ratio) became greater than the probability of producing overextended sequences (overshooting the required ratio), demonstrating a remarkable ability to accurately explore then exploit newly adaptive forms of action. Similarly, the speed-based efficiency of actions that achieved reward also clearly improved with training, although this was somewhat restricted, such that the speed of the first chunk was the only structural segment to adapt. Any new addition to this scaffold remained invariant, including the “joint” segment that melded the first scaffold sequence with *de novo* chunks. In this self-paced sequence task, adaptive responding was strictly dependent on an increasing fixed ratio schedule and not on target reward cues or time penalties, yet the consistency of action timing seen in later schedules indicates that skilled action timing is usefully transferred when updating contingencies. The high degree of internal cohesion (Mechner, 1958), both within and between the chunked units of action in a sequence, also suggests that the integration of chunks into whole sequences occurs smoothly, without disruption to the consistent timing of consecutive actions.

### M1→DLS circuitry and the injection of temporal variability to action

Considering the anatomical evidence and arguments supporting meaningful functional interactions between the motor cortical regions and diverse basal ganglia centres, and the effects on action timing we observed following M1-driven DLS lesion, we propose that the connectivity between motor cortical neurons and the DLS—perhaps through collaterals of descending pyramidal tract projections—may function to stabilize/destabilize the temporal boundaries of learned sequence durations by allocating the minimum level of variability required for explorative performance. The primary evidence for this view comes from the behavioural effects induced by specific ablation of the M1→DLS circuit. We found that this selective ablation did not alter the number of actions within successful or unsuccessful sequences *per se*, instead it induced briefer sequences with less varied durations that were ultimately less effective at earning reward (i.e., a slower reward rate and a greater proportion of unsuccessful actions). Perhaps unsurprisingly, we found that in the development of skilled actions, the timing of successful action sequences was more reliable (i.e. less variable) as task performance improved. However, when comparing the variability of unsuccessful actions relative to successful performance within-subjects, we showed that both Sham and M1→DLS lesion groups increased the variability of successful sequence durations, indicating that increased variation is a component of successful performance. Consistent with this argument is the observation that, in the less successful DLS lesion group, there was an overall reduced variance. The relationship between action timing variability and task success, therefore, may not be as simple as expecting variance to decline as task accuracy improves; rather, a broader baseline level of variance in action timing-space may be leveraged to expand the repertoire of available actions in action selection-space that is used for successful behavioural adaptation.

The effect of M1-driven DLS lesions on sequence duration was particularly pronounced in the first action chunk (presses 1-3), in which the brevity of sequence duration relative to controls was greatest in the initial exploratory phases of sequence acquisition, but diminished as the more successful sham animals sped-up their actions during training. The shortness and consistently reduced variability of sequence durations following the M1→DLS interrup-tion— and the waning yet ongoing M1→DLS input observed in other studies of motor skill behaviour (Kupferschmidt et al., 2017)—suggests that this circuitry modulates action sequence learning via its contribution to the degree of encoded variability (Neuringer, 2002) and its capacity to extend action sequence duration. Although in a different context, some prior evidence implicates subcortical processing (such as the hyperdirect pathway targeting the STN) in similar forms of temporal extension to optimise behaviour. One such example is the proposed ‘hold your horses’ effect, which provides the STN with the capacity to “buy time” when deliberating over difficult choices (Baunez et al., 2007; Baunez & Robbins, 1997; Frank, 2006; Frank et al., 2007). Here, in a similar way, subcortical circuitry is well placed to act as an online “noise injector”, providing the extra action time that is required for exploratory sequences to enhance their likelihood of success; perhaps directly through interactions with the M1→DLS projection or through broader bottom-up basal ganglia-thalamo-cortical processing. In either case, a greater range of possible action sequence durations, within which more varied action timing could fall, might be expected to facilitate the exploration process and the subsequent learning of the target sequence necessary for its exploitation—a kind of dual time/variability-based contribution to the exploration-exploitation tradeoff of action.

### Anatomy and motoric function of corticofugal accessory collaterals

In support of the above possibility, this study provided evidence of a direct interaction between lateral corticostriatal circuitry and the downstream basal ganglia network by mapping collaterals of long-range pyramidal tract projections ramifying into different stations of the basal ganglia, particularly the lateral territories of the striatum, the GPe and the STN. We ensured the shared origin of these collaterals by implementing a dual-viral tracing approach, by which M1 cortical neurons projecting to the STN were first labelled in isolation and then their synaptic terminals throughout the basal ganglia were subsequently mapped. This method identified collaterals to both the DLS and the GPe, with highest synaptic volumes in the DLS, extending previous research (Karube et al., 2019; Kita & Kita, 2012; Nelson et al., 2021). However, our method could not distinguish M1→STN projections with independent collaterals to DLS or GPe from collective M1→STN projections with collaterals to both. New trans-synaptic tracing technology able to disentangle this circuitry is forthcoming (Li et al., 2019). Nonetheless, the anatomical finding of shared connectivity suggests that the DLS and GPe may be responsible for processing some of the action timing information that has typically been attributed to exclusive collaterals to the STN—the so-called hyperdirect pathway. It is possible, however, that the local integration of this timing information in downstream structures may differ. For example, the local inter-cellular interactions between D2-SPNs and D1-SPNs within the striatum (Matamales et al., 2020) and between prototypic and arkypallidal cells within the GPe (Aristieta et al., 2021) have been implicated in adaptive learning and locomotion functions, respectively. The importance of these “upstream” targets of the accessory corticofugal projection aligns with similar descriptions of basal ganglia function described in the center-surround model (Nambu et al., 2002), in which information is processed in a feedback-loop that differentially recruits direct, indirect or hyperdirect pathways traversing “forward” and “backward” through the basal ganglia to control motor performance. Similarly, in rats, race models of basal ganglia-driven behavioural response inhibition describe competition between Go, Stop and Pause signals emerging from striatum, GPe and STN, respectively, from which the timing of each competing signal is integral to the eventual behavioural output (Logan & Cowan, 1984; Mallet et al., 2016; Schmidt & Berke, 2017). Presently, using models of the basal ganglia to predict the impact of multiple corticofugal collateral inputs within the circuity is speculative and requires further experimentation. For example, simultaneous *in vivo* recordings could demonstrate the temporal relationships of downstream firing that occurs in response to excitation from a motor cortical input ubiquitous to DLS, GPe and STN regions—further informing interpretations of race models. Just how the supply of shared motor cortical information to multiple basal ganglia nuclei governs function, both locally and at the circuit level, remains to be understood. Nevertheless, such broad projections suggest a widespread and coordinated integration of motor cortical efference copies, a process that is likely essential for adapting on-going streams of behaviour throughout learning. Overall, the anatomical mapping of shared striatofugal collaterals to various basal ganglia structures in conjunction with action timing and variability effects observed following M1→DLS lesion, provides an interesting new avenue for future experiments that connect the functional contribution of corticofugal-basal ganglia networks to action timing-based variability and its adaptive role in behavioural sequences of action.

## Acknowledgements

We thank Jennifer Strempel, Anne Rowan and Lydia Williams for assistance with animal care.

