## Supplementary Materials for "Dynamic adaptation of sequential action benefits from cortico-basal ganglia-related temporal variability"

www.NeuroModuLab.com

**This PDF file includes:**

Materials and Methods

Figure supplements

Figure 1 – figure supplement 1

Figure 1 – figure supplement 2

Figure 2 – figure supplement 1

Figure 3 – figure supplement 1

Figure 4 – figure supplement 1

Figure 4 – figure supplement 2

Table supplement 1

### Materials and Methods

#### SUBJECTS

All experimental procedures were approved by the Animal Care and Ethics Committee at the University of New South Wales in accordance with the Australian Code of Practice for the Care and Use of Animals for Scientific Purposes (National Health and Medical Research Council, 2013) and Animal Care and Ethics Committee (ACEC) guidelines. A total of 35 mice (C57Bl/6 and C57Bl/6 x Drd2-eGFP hybrids) including both male and females ranging from 3-6 months old were used in this study. Subjects for all experiments were mice (*Mus musculus*). Mouse home cages were stored in a climate-controlled colony room. Light/dark cycles rotated every 12 hours and cage air was ventilated with an air handling unit (Tecniplast, Italy). Cages were made of clear plastic and enriched with a blue plastic igloo, a red plastic cylinder and dried corn-based bedding. Mice were grouped into 2-6 littermates per home cage and had *ad libitum* access to water and standardised lab chow up to 2 days prior to the onset of the behavioural tasks.

Sixteen female and sixteen male DRD2-EGFP-F1 hybrid mice (RRID:MMRRC_000230-UNC, originally sourced from Jackson laboratory, US) were bred and maintained in-house and used in the behavioural observation study. Three male C57BL/6 mice (RRID:IMSR_ARC:B6; Animal Resource Center, AU) were included in the viral tracing study of accessory hyperdirect pathways. Sixteen male C57BL/6 mice were used in the study of motor cortex driven function of post-synaptic cells in the DLS. Any differences in the female-male sex ratios were due to limitations in availability.

#### BEHAVIOURAL PROCEDURES

##### Apparatus

All behavioural tasks were performed within operant conditioning chambers (Med Associates Inc, US) that were contained within light and sound attenuating cubicles. Each chamber was fitted with a pellet dispenser capable of delivering 20 mg grain pellets (#F0163, Bio-Serv, US) to a recessed feeding magazine, and two retractable levers separated by the magazine. An infrared photobeam spanned the entrance to the magazine, and any breaks in this photobeam were recorded as magazine entries. A light (3W, 24V) was situated at the top-center of the rear chamber wall, and was illuminated during behavioural experiments. Behavioural tasks were programmed using MedState notation (Pascal programming language) in order to control the extension and retraction of levers, delivery of pellets, and to turn the house light on and off. Programs were executed and behavioural responses (i.e. lever presses or magazine entries) and outcomes (i.e. pellet delivery) were detected and outputted to .txt files using Med-PC software (Med Associates Inc, US).

##### Food restriction

Animals' access to food was restricted from 2 days prior to and throughout the behavioural training period. Every 24hrs, following behavioural training, approximately 2.3 grams of standard lab chow per animal was left in their home cage. Mice weights and health indicators were measured and recorded each day, ensuring animals maintained a body weight above 85% of their free-feeding weight and a healthy disposition.

**Magazine training**

In all behavioural experiments, animals underwent one magazine training session per day for two days prior to instrumental training. Each mouse was assigned a chamber, which was sustained throughout the experiment. During magazine training, levers were retracted and 20 grain pellets (20 mg, 3.35 kcal/g each) were delivered to the magazine on a random-time 60-s schedule over 20 minutes.

**Tandem lever sequence task**

All instrumental training began with the insertion of the lever/s and the illumination of the chamber light and finished with the retraction of the lever/s and the extinction of the light. Each session occurred once a day, lasting up to 45 minutes or 20 pellet deliveries (whichever came first). In the 'tandem sequence task', instrumental training began with continuous reinforcement (CRF) on a single lever ('End lever') for 4 sessions, during which, each lever press resulted in the delivery of a grain pellet reward. In the following 4 training sessions, a second lever was extended in tandem with the lever that was previously rewarded. Animals were now required to press once on the newly introduced lever ('Sequence lever'), prior to pressing the previously rewarded lever ('End lever') in order to receive a reward. The number of presses required (fixed ratio - FR) on the Sequence lever increased by two presses every four sessions from FR1 to FR3 to FR5 to FR7 in the study that focussed on motor cortical driven lesion of the DLS. In the behavioural observation study, however, the required presses on the Sequence lever increased by two presses every four sessions from FR1 to FR3 to FR5, then sustained on FR5 for seven sessions to observe the influence of overtraining on well-established sequences. The left and right position of levers that served as either the Sequence or End lever were counterbalanced within each group.

**Behavioural analysis**

MedState notation scripts coded individual events as discrete numbers: Sequence lever press (1), End lever press (2), pellet delivery (3), and magazine entry (4). When any of these events occurred during training, the coded number was stored chronologically and timestamped (0.01s time resolution) in an output array and saved to a .txt file. MATLAB scripts extracted the raw data from these .txt files and performed data organisational functions, behavioural quantifications, and data representations. Data organisation comprised alignment of coded action numbers with timestamp data points, such that each action was indexed to its corresponding time of performance in the session. Data was further organised by coding event transition types. Here, all combinations of transitions between any of the four coded events (Sequence lever press, End lever press, pellet delivery, and magazine entry) were given unique numeric identifiers used to find their position in an action sequence. In addition to within-sequence analysis, sequences could be categorised based on their association with reward: sequences immediately followed by reward delivery were called '*Successful';* all sequences that did not result in reward were considered '*Unsuccessful';* and all sequences that occurred — irrespective of reward — were termed *'All'* sequences. Other more general measures of behaviour could also be extracted from this numerically action-coded dataset, including: totals and rates of Sequence lever and End lever presses, total reward deliveries and rates, and total magazine entries and rates. Analysis of the chronological arrangement of sequence chunks required indexing a specific numeric identifier within a sequence and calculating the time between this and the numeric identifier of interest. For example, when investigating the duration of Chunk 2 in FR5 training, the time from the 3$^{rd}$ press to the 5$^{th}$ press was calculated by subtracting the time at which the 5th press in the sequence occurred during the session from the time the 3rd press occurred in the session; the remained being the time between these two events. Each of these times were arranged chronologically for each mouse within a session and aligned between mice. This allowed for the collective comparison of chunk durations of all mice at any given chronological position to be compared against the chunk durations of all mice occurring at different chronological positions.

#### VIRAL PROCEDURES

##### Stereotaxic surgical injection of viruses

Animals were anaesthetised using isoflurane gas (Laser Animal Health, Pharmachem, AU). Induction commenced with 3% isoflurane delivered in oxygen at 0.5L/min in an induction chamber. After approximately five minutes animals were transferred to a stereotaxic frame (Kopf instruments) and fitted to the face mask and ear bars, and maintained on 1-1.5% isoflurane mixture with oxygen (0.5L/min). Fur above the cranium was removed with scissors and hair removal cream and the area was sanitised with betadine antiseptic solution. Bupivacaine (0.1 ml), a local anaesthetic, was subcutaneously administered at the surgical site, while Carprofen analgesic (0.4ml/kg) and saline (1ml) were delivered subcutaneously at the lower back. An incision was made on the scalp to reveal the skull, and animal head placement was adjusted to align bregma and lambda skull landmarks on the sagittal and transverse planes. Injection sites were determined by anterior-posterior, medial-lateral, and dorso-ventral (from skull) axis coordinates from "The Mouse Brain in stereotaxic coordinates, 3rd edition" (Paxinos & Franklin, 2007), and from pilot injection studies. Using a 26-gauge needle mounted to the stereotaxic holder, holes (~0.2mm width) through the skull were carefully pierced above injection sites relative to bregma. Infusion fluid (virus or control solution) was loaded into a microinjector (Nanoject III; Drummond Scientific Company), and its pulled glass capillary pipette tip (GC100TF-15; Harvard Apparatus) pulled using a micropipette puller (P-97, Sutter Instrument) and lowered through the puncture hole to the injection site. A 2-minute waiting period occurred prior to the injection, which was infused at a rate of 2 nl/second; total injection volumes varied depending on experiment (see below for details). Following the injection, infusion fluid was left to rest for three minutes before retracting the microinjector pipette tip. The incision site on the scalp was sutured with surgical thread and treated with Betadine Antiseptic Topical Ointment and sealed with tissue adhesive (3M Vetbond). The delivery of anaesthetic was stopped, and animals were laid on a heat mat for 5 minutes prior to placing them in a recovery cage.

##### Circuit-specific ablation of striatal neurons receiving motor cortical projections

To selectively ablate the post-synaptic targets of the motor cortical projections in the DLS, a combination of interacting viral systems was used. We stereotaxically injected a first AAV (500 nl) with anterograde transsynaptic profile expressing Cre (AAV2-EF1a-mCherry-IRES-WGA-Cre; Addgene #55632, RRID:Addgene_55632) (Fisher et al., 2020; Gradinaru et al., 2010; Yang et al., 2013) into the M1 region of the cortex (AP: -2.06 mm; ML: +1.58; DV : -4.9 (from skull)). To maximise the relevance of our manipulation to the hyperdirect pathway collaterals to the striatum, in our second viral injection we designed the coordinates based on the high terminal density zones defined in our previous viral tracing study (see Figure 4H and Figure 4–figure supplement 6A-C) (coordinate:

AP: 0 mm; ML: +2.65; DV: -3.0 [from skull]); volume (500-650 nl). Half of the animals received the AAV-Flex-taCasp3-TEVp virus (Addgene #45580, RRID:Addgene_45580), which induces expression of a designer procaspase 3 (taCasp3) that is lacking endogenous caspase cleavage sites but is sensitive to the heterologous tobacco etch virus protease (TEVp) in the presence of Cre (Group Lesion). The other half received, in the same region, an AAV5-EF1A-DIO-eYFP control virus expressing Cre-dependent eYFP (group Sham).

**Viral tracing of accessory hyperdirect pathway**

To assess if hyperdirect pathway from primary motor cortex (M1) to subthalamic nucleus (STN) also sends accessory projections from the same motor cortical neurons to the posterior dorsal striatum (pDStr) and/or to the external segment of the globus pallidus (GPe), a combination of two virus was injected intracranially into the STN and M1. First, 85 nl of retro-cre-EGFP (rAAV5-CMV-HI-EGFP-Cre-WPRE; Addgene #105545, RRID:Addgene_105545) was infused unilaterally into the STN (AP: -2.06 mm; ML: +1.58; DV: -4.9). Then, 400 nl of antero-mGFP-Syp-mRuby (AAV2-5-hSyn-FLEX-mGFP-2A-Synaptophysin-mRuby; Addgene #71760, RRID:Addgene_71760) was infused unilaterally into the M1 (AP: -1.78 mm; ML: +1.75; DV: -1.23) ipsilateral to the STN injection. This method allows for visualisation of both axonal projections (mGFP) and pre-synaptic boutons (Syp-mRuby) in anterograde synaptic territories, such as the pDStr, GPe and STN of Cre expressing motor cortical cells that are known to project to the STN.

#### TISSUE PROCESSING AND IMMUNOFLUORESCENCE LABELLING

**Transcardial fixation and tissue sectioning**

In behavioural experiments, animals were anesthetised with isoflurane gas (4% in air; Laser Animal Health, Pharmachem, AU) for 1 minute inside a sealed container. A lethal intraperitoneal injection of sodium pentobarbital (0.5-0.9ml, 500 mg/kg; Virbac Pty. Ltd., Australia) was administered, and follow-up paw and tail reflex checks occurred before commencing the perfusion. Mice were perfused transcardially for ten minutes using an air pressure system (15 ml/min flow) with 4% paraformaldehyde (PFA) in a solution of 0.1 M sodium phosphate buffer (pH 7.4). Brains were extracted and stored individually in PFA solution at 4°C for 12-48hrs before sectioning. Consecutive 30 µm coronal sections of brain were sliced in 0.1M phosphate buffer solution (PBS) using a vibratome (LEICA VT1000S, Leica Microsystems, Germany). Sections were sliced in an anterior-posterior direction, spanning regions of interest including: motor cortex, striatum, external globus pallidus and subthalamic nucleus. Free-floating sections were stored in a cryoprotectant solution (0.1 M sodium phosphate buffer, 30% v/v ethylene glycol, 30% v/v glycerol, 0.25 M Tris buffer) at -20°C until immunofluorescence procedures.

**Immunofluorescence labelling**

Free-floating sections were rinsed three times in Tris-buffered saline (TBS) solution (0.25 M Tris and 0.5 M NaCl at pH 7.4) for 10 min in an orbital shaker at room temperature. In the viral tracing experiment, sections were mounted on Superfrost Plus coated slides (Thermo Fisher Scientific) and Vectashield fluorescence medium (Vector Laboratories) was applied before placing a coverslip on top. The study that focussed on identifying a motor cortical driven lesion of the DLS required Nissl staining

(640/660 deep-red fluorescent Nissl Stain, cat. No. 21483; 1:500; Thermo Fisher Scientific, US) for cell quantification. Sections were washed three times for 10 minutes in TBS at room temperature before permeabilisation in 0.5% triton X-100 in TBS for 2hrs, followed by three 10-minute washes in TBS at room temperature. Sections were then incubated for 1hr at room temperature in Nissl stain, followed by three 10-minute washes in TBS at room temperature before mounting. Slides were stored at 4°C and images were captured within 72hrs of mounting.

#### IMAGE ACQUISITION AND QUANTITATIVE ANALYSIS

##### Spinning disk confocal microscopy

A wide-field spinning disk confocal system was used to capture images of mouse brain sections. A Diskovery multi-modal imaging platform and Zyla 4.2 sCMOS camera (Andor Technology) was added to a Nikon Eclipse TiE microscope body with a motorized stage and the Nikon Perfect Focus System, with image acquisition controlled by Nikon NIS-Elements software used to capture and produce mosaic images with 20x optical magnification, 16-bit pixel depth at 3.0269 pixels/µm image resolution. Up to three channels (488 nm, 561 nm and 640 nm lasers) were captured. An Olympus Confocal microscope (Olympus BX61WI) was also used to capture images at cortical injection sites, at 40x optical magnification at 3.2258 pixels/µm resolution and up to 2 channels (473 nm and 559 nm lasers) per image.

##### Nuclear and synaptic varicosity/bouton mapping

Spinning disk confocal images from each animal were processed using ImageJ2/Fiji software (v. 1.53c) (Schindelin et al., 2012). Freehand selections were used to create regional outlines of the DStr, GPe and STN and the regional area (mm2) and Cartesian (x,y) outline coordinates were measured. Binary images were generated from thresholds based on pixel intensity of biofluorescence in the soma (nissl fluorescence) or the synaptic boutons (Syp-mRuby). These binary images were used to quantify fluorescence with the *Analyze Particle* command, which locates the edge of an object based on its roundness and size, and determines the particles' cartesian (x,y) coordinates of its centroid position. Data on the position of the fluorescent particles and the regional outline were imported into MATLAB (MathWorks). The *inpolygon* function returned a list of points within the regional outline, and a distribution of the particles was reconstructed for each slice with a line-plot of the regional outline and scatter plot of cartesian centroid points. In the viral tracing study, the *densityplot* function was used to generate a colourmap relative to the spatial density of particles. Particle density was then calculated as the total number of particles within the regional outline area (particles/mm$^2$).

#### ALLEN MOUSE BRAIN CONNECTIVITY ATLAS RESOURCE METHODOLOGY

The Adult Mouse Connectivity Atlas is a large-scale searchable image database containing serial two-photon tomographic images of axonal projections labelled by viral (rAAV) tracers. The Atlas is built on an extensive library of experiments using enhanced green fluorescent protein (EGFP)-expressing adeno-associated viral vectors that are used to trace axonal projections from defined regions and cell types. These are imaged through high-throughput serial two-photon tomography to capture the EGFP-labelled axons throughout the brain. A computational model yields insights into connectional strength, distribution, symmetry and other network properties (Oh et al., 2014).

**Selection of Allen Mouse Brain Connectivity Atlas studies**

There are many approaches to searching the Adult Mouse Connectivity Atlas database, outlined here: http://help.brain-map.org/display/mouseconnectivity/Projection#Projection-Searching. In order to test the connectivity between the motor cortical regions and STN, we applied a specific filtered search to both the viral tracer injection site ('Source') and projection target structure ('Target'). Here we inputted both Primary (MOp) and Secondary (MOs) motor cortex as 'Source' structures and the STN as the 'Target' structure. Using the 'Source' and 'Target' search in this way we created a "virtual" anterograde study of the M2/M1 injection site and its projections to the STN. Searching was then filtered by projection density; calculated as a ratio of pixels with signal over all pixels in the structure. Three experiments with the highest projection density to the STN from cortical regions were selected for analysis. The details of these experiments can be found in Table 1.

**Table 1: Experimental details of studies from the Allen Mouse Brain Connectivity Atlas**

| **Experiment number** | **Mouse strain** | **Tracer type** | **Target region** | **Injection volume** (mm^3) | **Injection 1 coordinates** (mm from Bregma) | **Injection 2 coordinates** (mm from Bregma) |
|---|---|---|---|---|---|---|
| 180719293 | C57BL/6J | EGFP | anterior M2 region and dorsal agranalular insular area | 1.05 | A-P: +2.1<br>M-L: +2.4<br>D-V: -1<br>Angle: 30 | A-P: +2.1<br>M-L: +2.4<br>D-V: -0.4<br>Angle: 30 |
| 180709942 | C57BL/6J | EGFP | anterior M2/M1 region | 0.81 | A-P: +1.94<br>M-L: +2.5<br>D-V: -1.15<br>Angle: 0 | N/A |
| 100141780 | C57BL/6J | EGFP | anterior M1 region | 0.18 | A-P: +1.1<br>M-L: +2<br>D-V: -0.3<br>Angle: 20 | A-P: +1.1<br>M-L: +2<br>D-V: -0.75<br>Angle: 3 |

#### STATISTICAL ANALYSES

Statistical analysis was performed predominately using IBM SPSS Statistics software (version 28; IBM Corporation, NY). The a priori alpha level was set at $p < 0.05$. To test the assumption of equal variances between independent groups, Levene's test for equal variances was applied with the null hypothesis that the error variance was equal across groups. If the null hypothesis was rejected, Welch-Satterthwaite corrections to the degrees of freedom were applied (SPSS's 'equality of variance not assumed' option) to any independent samples t-tests (two-tailed). For univariate and repeated measures ANOVA, the homogeneity of variance was tested using Mauchly's test of sphericity. If the assumptions of sphericity were violated, Greenhouse-Geisser corrections were applied. Repeated measures ANOVA calculations require complete data; thus, in instances where a value was missing from a data set relative to another subject (e.g. when one mouse reached the maximum of 20 rewarded sequences in a session, while another did not), the fitting of a mixed effects model was used to analyse repeated measures data with missing values. A compound symmetry covariance matrix was used in the mixed effects model, and to control for assumptions of homogeneity of variance, Greenhouse-Geisser corrections were applied using Graphpad Prism (version 9; GraphPad Software, US). Linear regression analysis was also applied to infer the relationship between dependent variable/s and their occurrence chronologically.

#### DATA AVAILABILITY

All original raw data with statistical reports has been deposited at Figshare and will be made public as of the date of publication. All code used in this study will also be shared although is available upon request to the authors.

### Figure supplements

Figure 1 - figure supplement 1

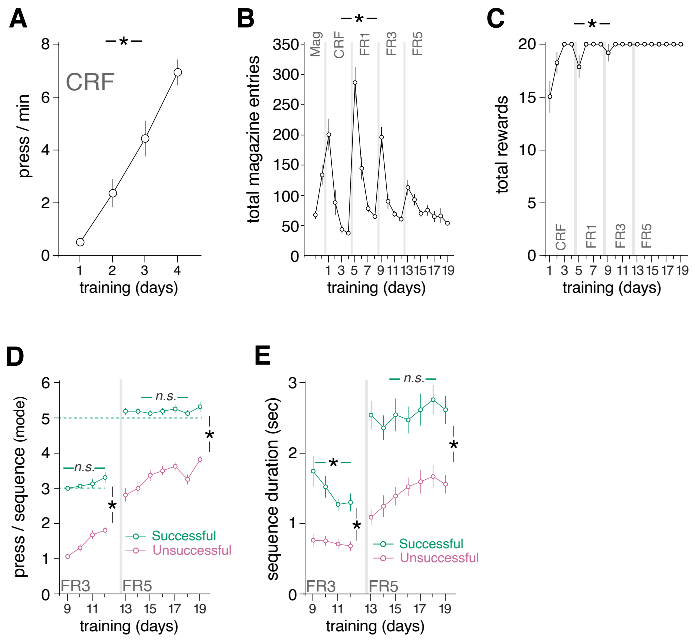


**Figure 1 - figure supplement 1. The tandem self-set sequence task generates instrumental performance measures and allows for distinction between successful and unsuccessful sequence data.** (**A**) Lever press rate (press/minute) during continuous reinforcement (CRF, End Lever) on Pre-Training (days 1-4). (**B**) Total magazine entries throughout all phases of training (Magazine training [Mag] → continuous reinforcement [CRF] → fixed ratio 1 [FR1] → fixed ratio 3 [FR3] → fixed ratio 5 [FR5]). (**C**) Total rewards earned during instrumental conditioning (CRF→FR1→FR3→FR5). (**D** and **E**) number of presses/sequence (D) and duration (E) of successful and unsuccessful sequences during FR3→FR5 training. *, significant overall/simple effect (black). N.S., not significant (Table supplement 1).

Figure 1 - figure supplement 2

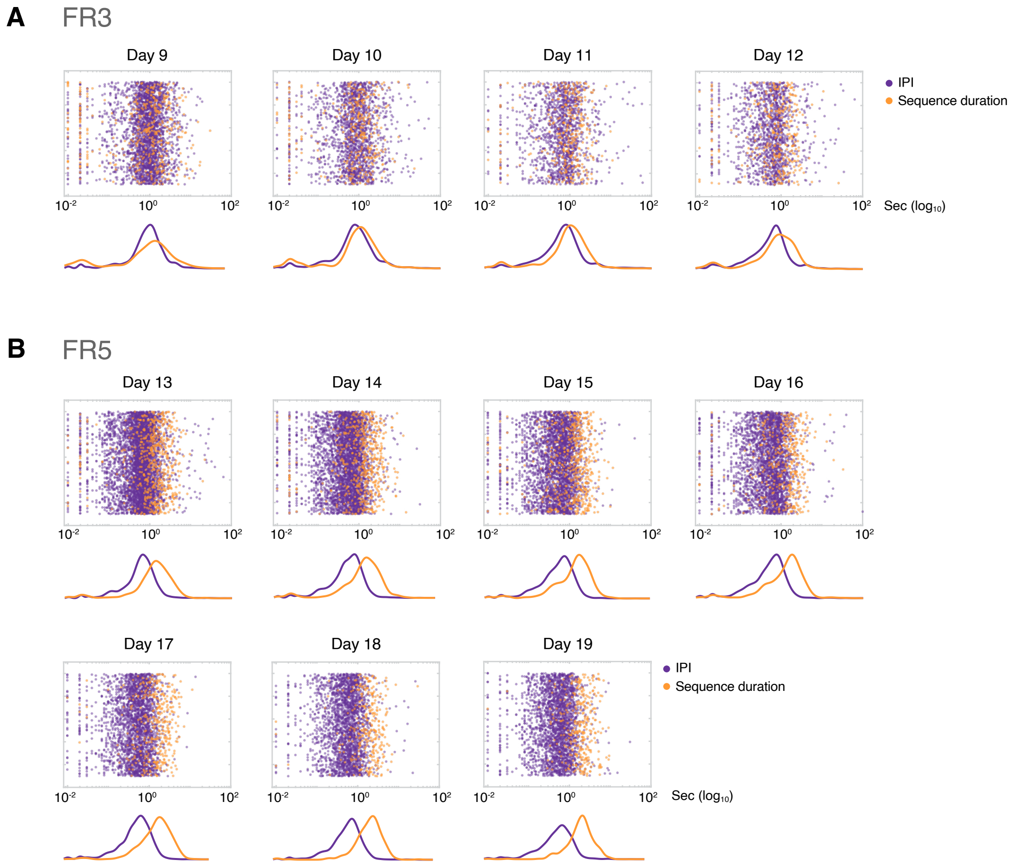


**Figure 1 - figure supplement 2. IPI and sequence duration functions dissociate throughout action sequence learning.** (**A** and **B**) Scatter plot of each IPI and sequence duration value (transformed to $log_{10}$) on the Sequence lever for all animals during FR3 (A; days 9-12) and FR5 (B; days 13-19), with corresponding fitted probability density function curves. Each dot corresponds to one IPI or one sequence duration from all mice of the experimental cohort. Data are distributed randomly in the Y axis.

Figure 2 - figure supplement 1

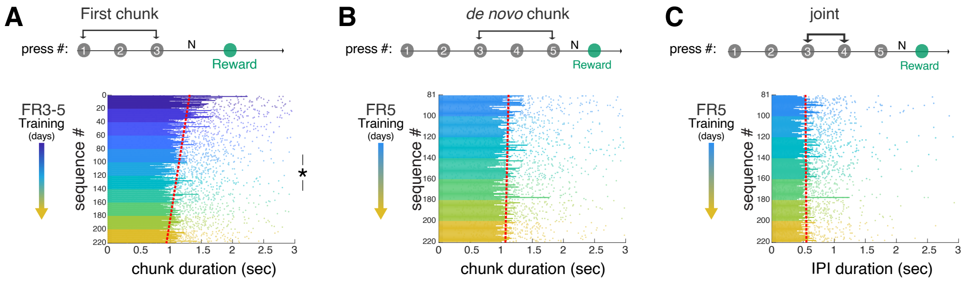


**Figure 2 - figure supplement 1. Only the first sequence prototypes evolve temporally during action sequence learning.** Duration of each successful sequence produced by all mice during training is plotted in chronological order across indicated phases of training (change in colour). Data are the duration of each sequence by each mouse (dots) and the average across mice (bars). A linear regression model highlighting the chronological trend is fitted to the data (red dashed line). (**A**) Duration of the very first successful subsequence segments (first chunk) throughout FR3→FR5. (**B**) Duration of the new part of the sequence (de novo chunk) added to the previous sequence scaffold (first chunk) in all successful sequences across the entire FR5 training. (**C**) Duration of the "joint" IPI (sits between the first and the de novo chunks) in all successful sequences across the entire FR5 training.

Figure 3 - figure supplement 1

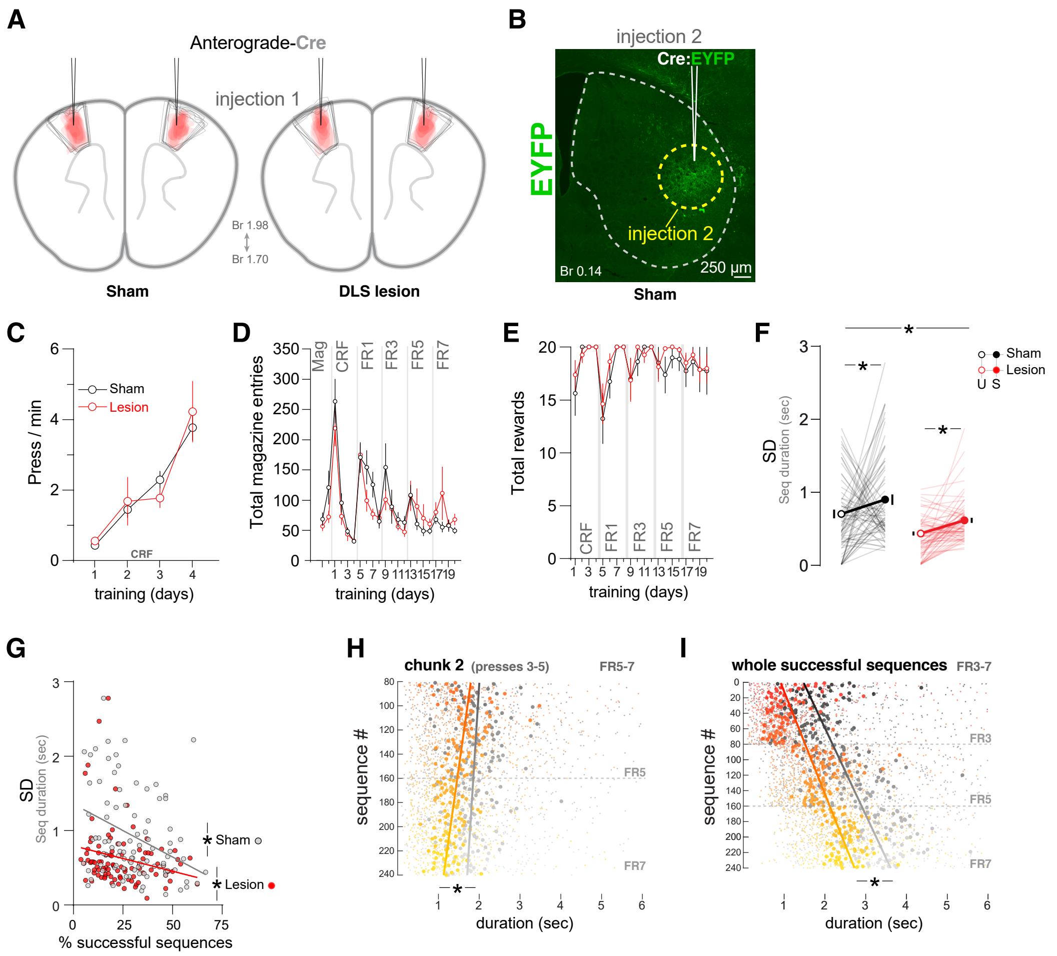


**Figure 3 - figure supplement 1. Injection sites and additional behavioural outcomes.** (**A**) Injection site of anterograde-cre (AAV2-EF1a-mCherry-IRES-WGA-Cre-Cre) in M1 for pre-synaptic driven lesion of post-synaptic DLS targets. Outline of injection site was overlayed for each animal in the Sham group (left) the DLS Lesion group (right); 1 slice/animal. (**B**) Example spinning disk confocal image (20x magnification) of Sham group animal that received Cre-dependent EYFP virus in the DLS high density zone. (**C**) Lever press rate (press/minute) during continuous reinforcement (CRF) for both Sham and DLS Lesion group. (**D**) Total magazine entries from magazine training through instrumental conditioning (CRF→FR1→FR3→FR5→FR7). (**E**) Total rewards during instrumental conditioning (CRF→FR1→FR3→FR5→FR7). (**F**) SD of successful (S) and unsuccessful (U) sequence durations (sec) for both Sham and DLS Lesion groups during combined sequence schedules (FR3+FR5+FR7). (**G**) SD of successful sequence durations (sec) as % successful sequences increase for both Sham and DLS Lesion during combined sequence schedules (FR3+FR5+FR7). (**H** and **I**) Duration of successful sub-sequence chunk 2 (press 3-5) during FR3→FR5 (H), and successful whole sequences (rewarded sequences above FR threshold) during FR1→FR3→FR5→FR7 (I), arranged chronologically for each mouse (small dots) and averaged across mice (large dots) for both Control and DLS lesion groups; with a linear regression model fitted to the data (red line). *, significant overall/simple effect (black) (Table supplement 1).

Figure 4 - figure supplement 1

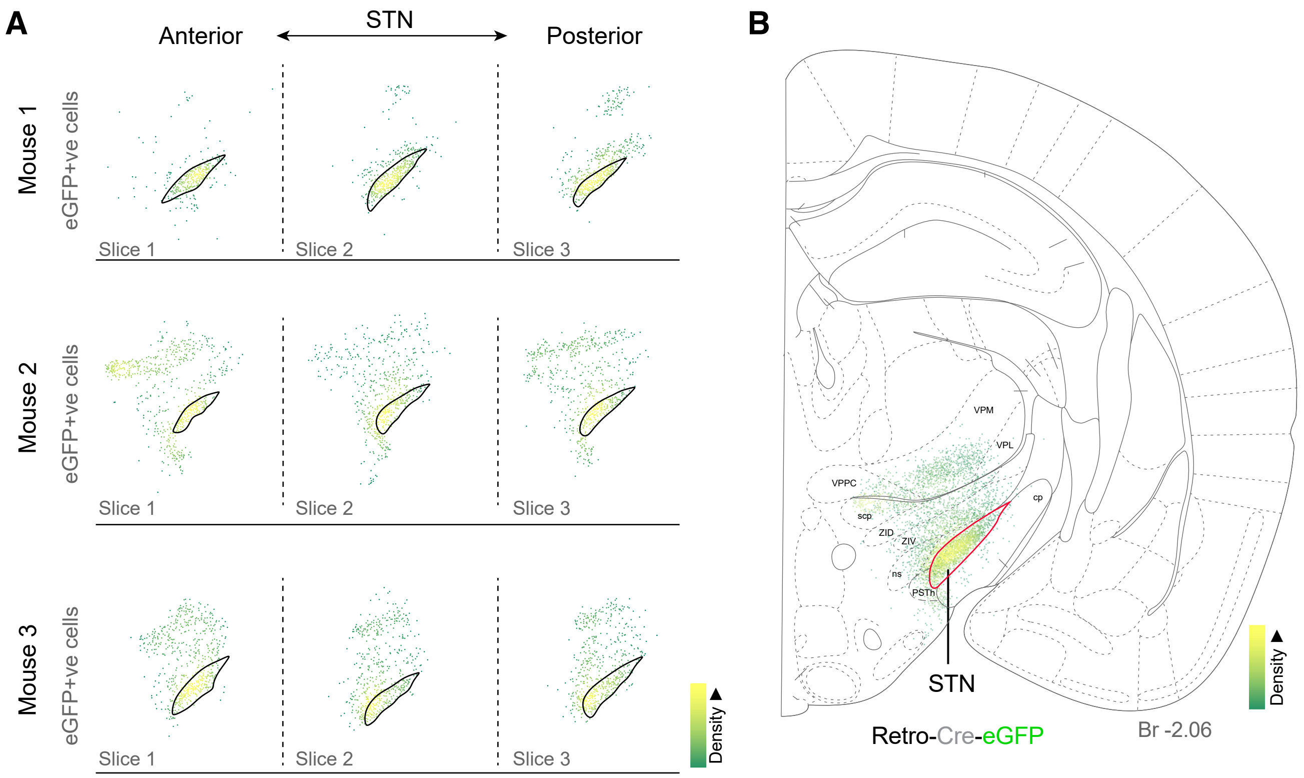


**Figure 4 – figure supplement 1. Density mapping of Retro-Cre-EGFP particles in the targeted STN.** (**A**) Reconstruction of eGFP particles after injection of Retro-Cre-eGFP virus in the STN. Three slices from 3 injected mice are represented, arranged from anterior to posterior. Each dot is an eGFP+ neuron. Density gradients are represented by a green-to-yellow pseudocoloured LUT palette (density maps). (**B**) Density maps overlaid for each slice (3x slice/animal; n = 3), fitted to adapted stereotaxic mouse atlas image at bregma -2.06 (Paxinos & Franklin, 2007).

Figure 4 - figure supplement 2

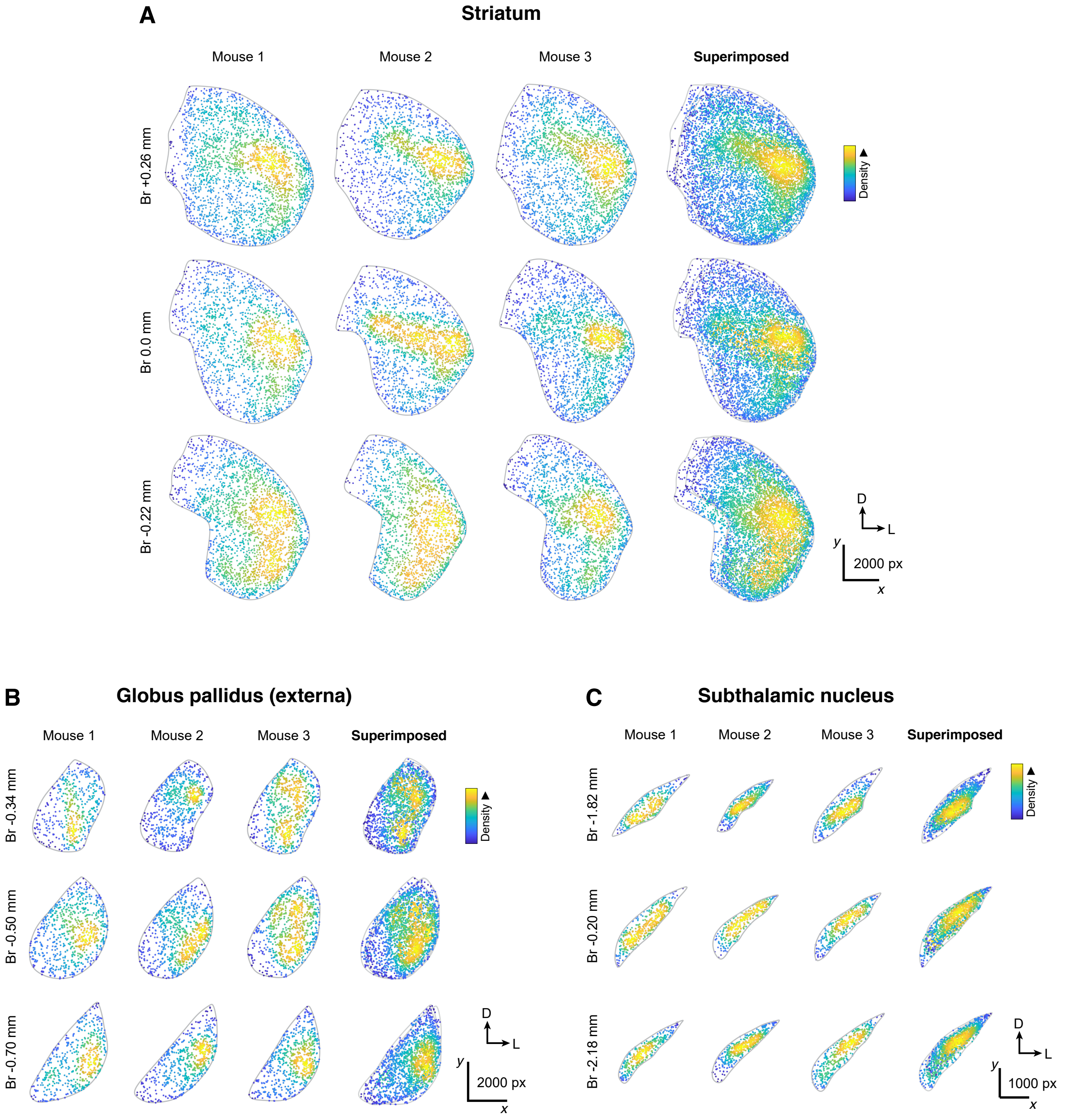


**Figure 4 – figure supplement 2. Pre-synaptic terminal distribution of cortico→STN collaterals in the striatum, GPe and target STN.** Mice received a first injection of Retro-Cre-eGFP in the STN (see Figure 4 – figure supplement 1) and a second injection of Antero-mGFP-Syp-mRuby in the M1 cortex (see Figure 4D and Methods). (**A-C**) Particle density maps of synaptophysin (Syp)-mRuby+ terminals in the striatum (A), GPe (B) and STN (C). All slices (3x slice/animal; n = 3) on the side ipsilateral to STN and M1 injection sites are displayed individually, and displayed superimposed according to their anterior-posterior position. Density gradients are represented by a blue-green-yellow pseudocoloured LUT palette (density maps).

### Table supplements

**Table supplement 1. Reported statistics.** The a priori alpha level was set at $p<0.05$. To test the assumption of equal variances between independent groups, Levene's test for equal variances was applied to test the null hypothesis that the error variance was equal across groups. If the null hypothesis was rejected, Welch-Satterthwaite corrections to the degrees of freedom were applied to any independent samples t-tests (two-tailed). For repeated measures ANOVA, homogeneity of variance was tested using Mauchly's test of sphericity. If the assumptions of sphericity were violated, Greenhouse-Geisser corrections were applied. A compound symmetry covariance matrix was used when fitting the mixed effects model to repeated measures data with missing values, and to control for assumptions of homogeneity of variance, Greenhouse-Geisser corrections were applied. Linear regression analysis was applied to infer the relationship between dependent variable/s and their chronological occurrence. FXfsx; Figure X – figure supplement x.

| Figure | Analysis | Test | Dependent Variable/s | Independent Variable/s | Statistics | P value |
|---|---|---|---|---|---|---|
| **1B** | Lever press rate for Sequence and End lever across all training (FR1-FR3-FR5) | Repeated measures Two-way mixed ANOVA | presses/ min | Session [FR1-FR5]<br>- *within-subjects*<br><br>Lever<br>[Sequence or End]<br>- *between subjects* | Mauchly's sphericity:<br><br>*simple effects* (session)<br>$F_{(4.170,125.088)} = 51.031$<br><br>*main effects* (lever)<br>$F_{(1,30)} = 31.457$<br><br>*interaction* (session*lever):<br>$F_{(4.170,125.088)} = 92.048$ | $p < 0.001$<br><br><br>$p < 0.001$<br><br><br>$p < 0.001$<br><br><br><br>$p < 0.001$ |
| **1C** | Presses per sequence for Sequence and End lever across sequence training (FR3-FR5) | Repeated measures Two-way mixed ANOVA | presses/ sequence | Session [FR3-FR5]<br>- *within-subjects*<br><br>Lever<br>[Sequence or End]<br>- *between subjects* | Mauchly's sphericity:<br><br>*simple effects* (session)<br>$F_{(5.418,162.541)} = 56.178$<br><br>*main effects* (lever)<br>$F_{(1,30)} = 389.212$<br><br>*interaction* (session*lever)<br>$F_{(5.418,162.541)} = 55.083$ | $p < 0.001$<br><br><br>$p < 0.001$<br><br><br>$p < 0.001$<br><br><br><br>$p < 0.001$ |
| **1D** | Pellets per sequence across training (FR1, FR3 and FR5) | Repeated measures One-way ANOVA | pellets/ minute | Session [FR1, FR3 or FR5]<br>- *within-subjects* | Mauchly's sphericity:<br>FR1<br>FR3<br>FR5<br><br>*simple effects* (session)<br>FR1: $F_{(3,45)} = 88.259$<br>FR3: $F_{(3,45)} = 30.613$<br>FR5: $F_{(2.929,43.932)} = 8.311$ | <br>$p > 0.05$<br>$p > 0.05$<br>$p < 0.05$<br><br><br>$p < 0.001$<br>$p < 0.001$<br>$p < 0.001$ |
| **1E** | Number of sequences across training (FR3 and FR5) | Repeated measures One-way ANOVA | # seq | Session [FR3 or FR5]<br>- *within-subjects* | Mauchly's sphericity:<br>FR3<br>FR5<br><br>*simple effects* (session)<br>FR3: $F_{(1.579,23.679)} = 19.239$<br>FR5: $F_{(1.845,27.681)} = 9.544$ | <br>$p < 0.001$<br>$p < 0.001$<br><br><br>$p < 0.001$<br>$p < 0.001$ |
| **1G** | Inter-press intervals (peak probability | Repeated measures | Inter-press intervals (sec) | Session [FR3, FR5 or FR3-FR5]<br>- *within-subjects* | Mauchly's sphericity:<br>FR3<br>FR5 | <br>$p < 0.05$<br>$p < 0.05$ |

| Figure | Analysis | Test | Dependent Variable/s | Independent Variable/s | Statistics | P value |
|---|---|---|---|---|---|---|
| | density) across training (FR3 and FR5). | One-way ANOVA | | | FR3-FR5<br><br>*simple effects* (session)<br>FR3: $F_{(2.047,30.708)} = 1.991$<br>FR5: $F_{(3.402,51.028)} = 0.894$<br>FR3-FR5: $F_{(4.548,68.226)} = 4.291$ | $p < 0.05$<br><br><br>$p > 0.05$<br>$p > 0.05$<br>$p < 0.05$ |
| **1G** | Sequence duration (peak probability density) across training (FR3 and FR5) | Repeated measures One-way ANOVA | Sequence duration (sec) | Session [FR3 or FR5]<br>- *within-subjects* | Mauchly's sphericity:<br>FR3<br>FR5<br><br>*simple effects* (session)<br>FR3: $F_{(1.038,15.567)} = 8.206$<br>FR5: $F_{(2.568,38.527)} = 12.573$ | <br>$p < 0.001$<br>$p < 0.05$<br><br><br>$p < 0.05$<br>$p < 0.001$ |
| **1H** | Percentage of successful sequences across training (FR3 and FR5) | Repeated measures One-way ANOVA | % Successful sequences | Session [FR3 or FR5]<br>- *within-subjects* | Mauchly's sphericity:<br>FR3<br>FR5<br><br>*simple effects* (session)<br>FR3: $F_{(3,45)} = 24.223$<br>FR5: $F_{(6,90)} = 18.669$ | <br>$p > 0.05$<br>$p > 0.05$<br><br><br>$p < 0.001$<br>$p < 0.001$ |
| **1I** | Sequence duration (PPD) for successful & unsuccessful sequences during training (FR3 and FR5) | Paired two-tailed t-test | Sequence duration (sec) | Sequence type [successful or unsuccessful] | *Paired samples test:*<br><br>FR3: $T_{(63)} = 10.893$<br>FR5: $T_{(111)} = 21.377$ | <br><br>$p < 0.001$<br>$p < 0.001$ |
| **2A** | Presses per sequence (successful sequences) across sequence training (FR3 and FR5) | Repeated measures one-way ANOVA | presses/ sequence (successful sequences) | Session [FR3 or FR5]<br>- *within-subjects* | Mauchly's sphericity:<br>FR3<br>FR5<br>*simple effects* (session)<br>FR3: $F_{(1.781,26.715)} = 2.333$<br>FR5: $F_{(3.430,51.445)} = 0.556$ | <br>$p < 0.05$<br>$p < 0.05$<br><br>$p > 0.05$<br>$p > 0.05$ |
| **2A** | Presses per sequence of unsuccessful compared to successful sequences across sequence training (FR3 and FR5) | Repeated measures Two-way mixed ANOVA | presses/ sequence | Session [FR3 or FR5]<br>- *within-subjects*<br><br>Sequence type [unsuccessful or successful]<br>- *between subjects* | Mauchly's sphericity:<br>FR3<br>FR5<br><br>*Interaction* (session*lever)<br>FR3: $F_{(3,90)} = 3.220$<br>FR5: $F_{(3,90)} = 3.438$ | <br>$p > 0.05$<br>$p < 0.05$<br><br><br>$p < 0.001$<br>$p < 0.05$ |
| **2B (left)** | Frequency of certain sequence length (presses/ sequence) during FR3 | Total number (frequency ) | Frequency of #presses/ sequence | #presses/sequence [unsuccessful & successful]<br><br>*Unsuccessful*<br>(Presses 1-2)<br><br><br><br><br>*Successful*<br>(Presses 3-13) | *Frequency*<br><br>Presses/seq Frequency<br>1 2116<br>2 1460<br>3 726<br>4 321<br>5 131<br>6 47<br>7 20<br>8 8<br>9 6<br>10 4<br>11 2<br>12 1<br>13 1 | |
| **2B (left) inset** | Probability of performing a sequence of a certain length (presses/ sequence) during FR3 | Probability density function | Frequency of #presses/ sequence | #presses/sequence [unsuccessful & successful]<br><br>*Unsuccessful*<br>(Presses 1-2)<br><br><br><br><br>*Successful*:<br>(Presses 3-13) | *Probability density function*<br><br>Presses/seq Probability<br>1 0.591<br>2 0.408<br>3 0.573<br>4 0.253<br>5 0.103<br>6 0.037<br>7 0.016 | |

| Figure | Analysis | Test | Dependent Variable/s | Independent Variable/s | Statistics | P value |
|---|---|---|---|---|---|---|
| | | | | | 8 0.006<br>9 0.005<br>10 0.003<br>11 0.002<br>12 0.001<br>13 0.001 | |
| **2B (right)** | Frequency of certain sequence length (presses/ sequence) during FR5 | Total number (frequency) | Frequency of #presses/ sequence | #presses/sequence [unsuccessful & successful]<br><br>*Unsuccessful*<br>(Presses 1-4)<br><br>*Successful*<br>(Presses 5-18) | *Probability density function*<br>Presses/seq Frequency<br>1 442<br>2 1368<br>3 1741<br>4 1478<br>5 1010<br>6 597<br>7 309<br>8 146<br>9 86<br>10 45<br>11 19<br>12 11<br>13 2<br>14 7<br>15 2<br>16 2<br>17 1<br>18 2 | |
| **2B (right) inset** | Probability of performing a sequence of a certain length (presses/ sequence) during FR5 | Probability density function | Frequency of #presses/ sequence | #presses/sequence [unsuccessful & successful]<br><br>*Unsuccessful*<br>(Presses 1-4)<br><br>*Successful*<br>(Presses 5-18) | *Probability density function*<br><br>Presses/seq Probability<br>1 0.088<br>2 0.272<br>3 0.346<br>4 0.294<br>5 0.451<br>6 0.267<br>7 0.138<br>8 0.065<br>9 0.038<br>10 0.020<br>11 0.008<br>12 0.005<br>13 0.001<br>14 0.003<br>15 0.001<br>16 0.001<br>17 0.000<br>18 0.001 | |
| **2C** | Chronological change in successful sequence duration of the first chunk (press 1-3) across all FR3 sessions | Simple linear regression | Chronological Sequence duration (sec) | sequence # | *linear regression*<br>$F_{(1,78)} = 40.221$; $R^2 = 0.340$ | $p < 0.001$ |
| **2D** | Chronological change in successful sequence duration of the second chunk (press 3-5) across first 4 sessions of FR5 | Simple linear regression | Chronological Sequence duration (sec) | sequence # | *linear regression*<br>$F_{(1,78)} = 1.055$; $R^2 = 0.013$ | $p > 0.05$ |
| **2E** | Chronological change in successful sequence Inter-chunk interval (IPI 3-4) across first 4 sessions of FR5 | Simple linear regression | Chronological IPI (sec) | sequence # | *linear regression*<br>$F_{(1,78)} = 0.620$; $R^2 = 0.008$ | $p > 0.05$ |
| **2F** | Comparisons of individual inter-press intervals (peak probability density) within successful | Univariate ANOVA | Inter-press intervals (IPI) (sec) | IPI position<br>FR3: [IPI 1-2 and IPI 2-3]<br>- *between subjects* | *IPI position*<br>FR3: $F_{(1,126)} = 2.184$<br><br>*IPI position* | $p > 0.05$ |

| Figure | Analysis | Test | Dependent Variable/s | Independent Variable/s | Statistics | P value |
|---|---|---|---|---|---|---|
| | sequences across training (FR3 and FR5). | | | FR3: [IPI 1-2, IPI 2-3, IPI 3-4, IPI 4-5]<br>*- between subjects* | FR5: $F_{(1,444)}$ = 1.865 | p > 0.05 |
| **3B** | Density of Nissl +ve cells in the DLS high density region | two-tailed independent samples t-test | Nissl cells/mm$^2$ | group<br>[Sham and DLS lesion]<br>*- between subjects* | *Levene's test*<br><br>*main effect* (group)<br>$T_{(94)}$ = 4.326 | p > 0.05<br><br><br>p < 0.001 |
| **3C (left)** | Lever press rate for the Sequence and End lever for *Sham* animals across sequence training (FR3-FR5-FR7) | Repeated measures Two-way mixed ANOVA | presses/min | session [FR3-FR7]<br>*- within-subjects*<br><br>lever<br>[Sequence or End]<br>*- between subjects* | Mauchly's sphericity:<br><br><br>*interaction*<br>(session*lever): $F_{(3.950,55.305)}$ = 11.367 | p < 0.001<br><br><br><br>p < 0.001 |
| **3C (right)** | Lever press rate for the Sequence and End lever for *DLS lesion* animals across sequence training (FR3-FR5-FR7) | Repeated measures Two-way mixed ANOVA | presses/min | session [FR3-FR7]<br>*- within-subjects*<br><br>lever<br>[Sequence or End]<br>*- between subjects* | Mauchly's sphericity:<br><br><br>*interaction*<br>(session*lever): $F_{(2.636,29.229)}$ = 11.744 | p < 0.001<br><br><br><br>p < 0.001 |
| **3D (right)** | Reward rate for Sham and DLS lesion animals combined across sequence training (FR3-FR5-FR7) | two-tailed indepen-dent samples t-test | pellets/minute | group<br>[Sham and DLS lesion]<br>*- between subjects* | *Levene's test*<br><br>*main effect* (group)<br>$T_{(190)}$ = 2.273 | p > 0.05<br><br><br>p < 0.05 |
| **3E (right)** | Percentage of successful sequences for Sham and DLS lesion animals combined across sequence training (FR3-FR5-FR7) | two-tailed indepen-dent samples t-test | % successful (rewarded) sequences | group<br>[Sham and DLS lesion]<br>*- between subjects* | *Levene's test*<br><br>*main effect* (group)<br>$T_{(190)}$ = 2.926 | p > 0.05<br><br><br>p < 0.05 |
| **3F** | Presses per sequence of unsuccessful compared to successful sequences in Sham and DLS lesion group animals during sequence training (FR3, FR5, or FR7) | Repeated measures Two-way mixed ANOVA | presses/sequence (mode) | sequence type<br>[unsuccessful or successful]<br>*- within-subjects*<br><br>group<br>[Sham or DLS lesion]<br>*- between subjects* | *simple effect* (sequence type)<br>FR3: $F_{(1,62)}$ = 478.474<br>FR5: $F_{(1,62)}$ = 418.118<br>FR7: $F_{(1,62)}$ = 344.050<br><br>*main effect* (group)<br>FR3: $F_{(1,62)}$ = 2.896<br>FR5: $F_{(1,62)}$ = 3.225<br>FR7: $F_{(1,62)}$ = 3.225<br><br>*interaction*<br>(sequence type*group)<br>FR3: $F_{(1,62)}$ = 0.049<br>FR5: $F_{(1,62)}$ = 1.222<br>FR7: $F_{(1,62)}$ = 2.766 | <br>p < 0.001<br>p < 0.001<br>p < 0.001<br><br><br>p > 0.05<br>p > 0.05<br>p > 0.05<br><br><br><br>p > 0.05<br>p > 0.05<br>p > 0.05 |
| **3G** | Duration of unsuccessful compared to successful sequences in Sham and DLS lesion group animals during sequence training (FR3, FR5, or FR7) | Repeated measures Two-way mixed ANOVA | Sequence duration [peak probability density (seconds)] | sequence type<br>[unsuccessful or successful]<br>*- within-subjects*<br><br>group<br>[Sham or DLS lesion]<br>*- between subjects* | *simple effect* (sequence type)<br>FR3: $F_{(1,62)}$ = 150.691<br>FR5: $F_{(1,62)}$ = 141.563<br>FR7: $F_{(1,62)}$ = 92.437<br><br>*main effect* (group)<br>FR3: $F_{(1,62)}$ = 15.510<br>FR5: $F_{(1,62)}$ = F(1,62) = 8.773<br>FR7: $F_{(1,62)}$ = 11.541<br><br>*interaction*<br>(sequence type*group)<br>FR3: $F_{(1,62)}$ = 12.755<br>FR5: $F_{(1,62)}$ = 5.129<br>FR7: $F_{(1,62)}$ = 6.152<br><br>*Pairwise comp. (Bonferroni)*<br>Unsuccessful sequences: | p < 0.001<br>p < 0.001<br>p < 0.001<br><br><br>p < 0.001<br>p < 0.05<br>p < 0.05<br><br><br><br>p < 0.05<br>p < 0.05<br>p < 0.05<br><br><br><br>p < 0.05 |

| Figure | Analysis | Test | Dependent Variable/s | Independent Variable/s | Statistics | P value |
|---|---|---|---|---|---|---|
| | | | | | FR3,5 or 7: Sham > DLS lesion<br><br>Successful sequences:<br>FR3,5 or 7: Sham > DLS lesion | <br><br>$p < 0.05$ |
| **3H (top row)** | SD of unsuccessful sequence duration for Sham and DLS lesion animals in sequence training (FR3, FR5 or FR7) | two-tailed indepen-dent samples t-test | Standard deviation (SD) | group<br>[Sham and DLS lesion]<br>- *between subjects* | *Levene's test*<br>FR3:<br>FR5:<br>FR7:<br><br>*main effect* (group)<br>FR3: $T_{(51.488)} = 2.113$<br>FR5: $T_{(41.640)} = 3.005$<br>FR7: $T_{(36.297)} = 2.707$ | <br>$P < 0.05$<br>$P < 0.001$<br>$P < 0.001$<br><br><br>$p < 0.05$<br>$p < 0.05$<br>$p < 0.05$ |
| **3H (bottom row)** | SD of successful sequence duration for Sham and DLS lesion animals in sequence training (FR3, FR5 or FR7) | two-tailed indepen-dent samples t-test | Standard deviation (SD) | group<br>[Sham and DLS lesion]<br>- *between subjects* | *Levene's test*<br>FR3:<br>FR5:<br>FR7:<br><br>*main effect* (group)<br>FR3: $T_{(62)} = 1.623$<br>FR5: $T_{(50.039)} = 2.069$<br>FR7: $T_{(39.286)} = 3.038$ | <br>$p > 0.05$<br>$p < 0.05$<br>$p < 0.001$<br><br><br>$p > 0.05$<br>$p < 0.05$<br>$p < 0.05$ |
| **3I (left)** | Inter-press interval (IPI) of successful sequences in Sham and DLS lesion group animals during sequence training (FR3, FR5, or FR7) | Univariate ANOVA | Inter-press interval [peak probability density (seconds)] | IPI position<br>FR3: [1-2,2-3]<br>FR5: [1-2,2-3,3-4,4-5]<br>FR7: [1-2,2-3,3-4,4-5,5-6,6-7]<br>- *between-subjects*<br><br>group<br>[Sham or DLS lesion]<br>- *between subjects* | *main effect* (IPI position)<br>FR3: $F_{(1,124)} = 7.013$<br>FR5: $F_{(3,248)} = 1.663$<br>FR7: $F_{(5,372)} = 1.216$<br><br>*main effect* (group)<br>FR3: $F_{(1,124)} = 12.208$<br>FR5: $F_{(1,248)} = 19.778$<br>FR7: $F_{(1,372)} = 43.077$<br><br>*interaction*<br>(sequence type*group)<br>FR3: $F_{(1,124)} = 0.116$<br>FR5: $F_{(3,248)} = 0.762$<br>FR7: $F_{(5,372)} = 0.667$ | <br>$p < 0.05$<br>$p > 0.05$<br>$p > 0.05$<br><br><br>$p < 0.001$<br>$p < 0.001$<br>$p < 0.001$<br><br><br><br>$p > 0.05$<br>$p > 0.05$<br>$p > 0.05$ |
| **4G** | Total synaptophysin-mRuby (syp-mRuby) boutons in the DStr, GPe and STN | one-way ANOVA | Total syp-mRuby particles /region | region<br>[DStr, GPe and STN]<br>- *between subjects* | *main effect* (region)<br>$F_{(2,24)} = 226.840$<br><br>*Post-hoc (Bonferroni)*<br>*DStr > GPe or STN*<br>*GPe > STN* | <br>$p < 0.001$<br><br><br>$p < 0.001$<br>$p < 0.05$ |
| **4G** | Density of synaptophysin-mRuby (syp-mRuby) boutons in the DStr, GPe and STN | one-way ANOVA | syp-mRuby particles /$mm^2$ | region<br>[DStr, GPe and STN]<br>- *between subjects* | *main effect* (region)<br>$F_{(2,24)} = 105.098$<br><br>*post-hoc (Bonferroni)*<br>*STN > DStr or GPe*<br>*DStr > GPe* | <br>$p < 0.001$<br><br><br>$p < 0.001$<br>$p > 0.05$ |
| **F1-FS1A** | Lever press rate for Sequence lever across CRF training | Repeated measures one-way ANOVA | presses/ min | Session [CRF]<br>- *within-subjects* | Mauchly's sphericity:<br><br>*simple effects* (session)<br>CRF: $F_{(3,45)} = 70.689$ | $p > 0.05$<br><br><br>$p < 0.001$ |
| **F1-FS1B** | Total magazine entries across Magazine, CRF, FR1, FR3, FR5 training | Repeated measures one-way ANOVA | Total magazine entries | Session<br>- *within-subjects* | Mauchly's sphericity:<br><br>*simple effects* (session)<br>$F_{(20,300)} = 22.914$ | $p > 0.05$<br><br><br>$p < 0.001$ |
| **F1-FS1C** | Total rewards across CRF, FR1, FR3, FR5 training | Repeated measures one-way ANOVA | Total rewards | Session [CRF]<br>- *within-subjects* | Mauchly's sphericity:<br><br>*simple effects* (session)<br>$F_{(18,270)} = 5.797$ | $p > 0.05$<br><br><br>$p < 0.001$ |

| Figure | Analysis | Test | Dependent Variable/s | Independent Variable/s | Statistics | P value |
|---|---|---|---|---|---|---|
| **F1-FS1D** | Presses per sequence for Successful sequences across sequence training (FR3 or FR5) | Repeated measures one-way ANOVA | presses/ min | Session [FR3 or FR5] - *within-subjects* | FR3: Mauchly's sphericity<br>FR5: Mauchly's sphericity<br><br>*simple effects* (session)<br>FR3: $F_{(1.781,26.715)} =$ 2.333<br>FR5: $F_{(3.430,51.445)} =$ 0.556 | $p < 0.05$<br>$p < 0.05$<br><br><br>$p > 0.05$<br>$p > 0.05$ |
| **F1-FS1D** | Presses per sequence for Unsuccessful and Successful sequences across sequence training (FR3 or FR5) | Repeated measures Two-way mixed ANOVA | presses/ sequence (mode) | Session [FR3 or FR5] - *within-subjects*<br><br>Sequence type [unsuccessful or successful] - *between subjects* | FR3: Mauchly's sphericity<br>FR5: Mauchly's sphericity<br><br>*simple effect* (session)<br>FR3: $F_{(3,90)} =$ 13.272<br>FR5: $F_{(4.450,133.498)} =$ 5.282<br><br>*main effect* (sequence type)<br>FR3: $F_{(1,30)} = 312.111$<br>FR5: $F_{(1,30)} = 345.191$<br><br>*Interaction* (sequence type* session)<br>FR3: $F_{(3,90)} = 3.220$<br>FR5: $F_{(4.450,133.498)} = 3.438$ | $p > 0.05$<br>$p < 0.05$<br><br><br>$p < 0.001$<br>$p < 0.001$<br><br><br>$p < 0.001$<br>$p < 0.001$<br><br><br>$p < 0.001$<br>$p < 0.05$ |
| **F1-FS1E** | Sequence duration for Successful sequences across sequence training (FR3 or FR5) | Repeated measures one-way ANOVA | sequence duration [peak probability density (seconds)] | Session [FR3 or FR5] - *within-subjects* | FR3: Mauchly's sphericity<br>FR5: Mauchly's sphericity<br><br>*simple effects* (session)<br>FR3: $F_{(1.457,21.857)} =$ 5.584<br>FR5: $F_{(6,90)} =$ 1.151 | $p < 0.05$<br>$p < 0.05$<br><br><br>$p < 0.05$<br>$p > 0.05$ |
| **F1-FS1E** | Sequence duration for Unsuccessful and Successful sequences across sequence training (FR3 or FR5) | Repeated measures Two-way mixed ANOVA | sequence duration [peak probability density (seconds)] | Session [FR3 or FR5] - *within-subjects*<br><br>Sequence type [unsuccessful or successful] - *between subjects* | FR3: Mauchly's sphericity<br>FR5: Mauchly's sphericity<br><br>*simple effect* (session)<br>FR3: $F_{(1.617,48.520)} =$ 6.263<br>FR5: $F_{(4.054,121.607)} =$ 4.475<br><br>*main effect* (sequence type)<br>FR3: $F_{(1,30)} = 25.366$<br>FR5: $F_{(1,30)} = 28.026$<br><br>*Interaction* (sequence type* session)<br>FR3: $F_{(1.617,48.520)} = 3.393$<br>FR5: $F_{(4.054,121.607)} = 1.214$ | $p < 0.05$<br>$p < 0.05$<br><br><br>$p < 0.05$<br>$p < 0.001$<br><br><br>$p < 0.001$<br>$p < 0.001$<br><br><br>$p > 0.05$<br>$p > 0.05$ |
| **F2-FS1A** | Chronological change in successful sequence duration of the first chunk (press 1-3) across all FR3 & FR5 sessions | Simple linear regression | Chronological Sequence duration (sec) | sequence # | *linear regression*<br>$F_{(1,138)} = 74.081$; $R^2 = 0.254$ | <br>$p < 0.001$ |
| **F2-FS1B** | Chronological change in successful sequence duration of the first chunk (press 3-5) across all FR5 sessions | Simple linear regression | Chronological Sequence duration (sec) | sequence # | *linear regression*<br>$F_{(1,138)} = 1.485$; $R^2 = 0.011$ | <br>$p > 0.05$ |
| **F2-FS1C** | Chronological change in successful sequence inter-chunk interval (IPI 3-4) across all FR5 sessions | Simple linear regression | Chronological IPI (sec) | sequence # | *linear regression*<br>$F_{(1,138)} = 0.195$; $R^2 = 0.001$ | <br>$p > 0.05$ |

| Figure | Analysis | Test | Dependent Variable/s | Independent Variable/s | Statistics | P value |
|---|---|---|---|---|---|---|
| **F3-FS1C** | Lever press rate for the End lever across CRF | Repeated measures Two-way mixed ANOVA | presses/ min | session<br>- *within-subjects*<br><br>group<br>[Sham and DLS lesion]<br>- *between subjects* | Mauchly's sphericity:<br><br>*simple effect* (session)<br>$F_{(1.838,25.735)} = 27.307$<br><br>*main effect* (group)<br>$F_{(1,14)} = 0.040$<br><br>*interaction*<br>(session*group):<br>$F_{(1.838,25.735)} = 0.558$ | $p < 0.01$<br><br>$p < 0.001$<br><br>$p > 0.05$<br><br>$p > 0.05$ |
| **F3-FS1D** | Total number of magazine entries for *RET and EXT* groups across all training (magazine-CRF-FR1-FR3-FR5-FR7) | Repeated measures Two-way mixed ANOVA | Total magazine entries /session | session<br>- *within-subjects*<br><br>group<br>[Sham and DLS lesion]<br>- *between subjects* | Mauchly's sphericity:<br><br>*simple effect* (session)<br>$F_{(21,294)} = 15.563$<br><br>*main effect* (group)<br>$F_{(1,14)} = 0.343$<br><br>*interaction*<br>(session*group):<br>$F_{(21,294)} = 1.494$ | $p < 0.01$<br><br>$p < 0.001$<br><br>$p > 0.05$<br><br>$p > 0.05$ |
| **F3-FS1E** | Total number of rewards for *RET and EXT* groups across all training (magazine-CRF-FR1-FR3-FR5-FR7) | Repeated measures Two-way mixed ANOVA | presses/ min | session<br>- *within-subjects*<br><br>group<br>[Sham and DLS lesion]<br>- *between subjects* | Mauchly's sphericity:<br><br>*simple effect* (session)<br>$F_{(21,294)} = 4.154$<br><br>*main effect* (group)<br>$F_{(1,14)} = 0.361$<br><br>*interaction*<br>(session*group):<br>$F_{(21,294)} = 0.353$ | $p < 0.01$<br><br>$p < 0.001$<br><br>$p > 0.05$<br><br>$p > 0.05$ |
| **F3-FS1F** | SD of sequence duration in Unsuccessful and Successful sequences for Sham and DLS lesion groups across all sequence training (FR3-FR5-FR7) | Repeated measures Two-way mixed ANOVA | SD sequence duration (seconds) | sequence type [unsuccessful or successful]<br>- *within-subjects*<br><br>group<br>[Sham and DLS lesion]<br>- *between subjects* | *simple effect* (sequence type)<br>$F_{(1,190)} = 18.576$<br><br>*main effect* (group)<br>$F_{(1,190)} = 21.894$<br><br>*Interaction* (sequence type* group)<br>$F_{(1,190)} = 2.398$ | $p < 0.001$<br><br>$p < 0.001$<br><br>$p > 0.05$ |
| **F3-FS1G** | Percentage of successful sequences as SD of successful sequence duration for Sham and DLS lesion animals in sequence training (FR3-FR5-FR7) | Simple linear regression<br><br>one-way ANCOVA | Percentage of successful sequences | SD of successful sequence duration<br><br>group<br>[Sham and DLS lesion]<br>- *between subjects* | *linear regression (Sham)*<br>$F_{(1,94)} = 11.018$; $R^2 = 0.105$<br><br>*linear regression (DLS lesion)*<br>$F_{(1,94)} = 5.061$; $R^2 = 0.051$<br><br>*main effect* (group)<br>F(1,189)=15.946, p<0.001<br><br>*Regression parameter estimate(group)*<br>$T_{(2.074)}$=3.993, β-coefficient = -8.284 | $p < 0.01$<br><br>$p < 0.05$<br><br>$p < 0.001$<br><br>$p < 0.001$ |
| **F3-FS1H** | Duration of chunk 2 of successful sequences as they occur chronologically in Sham and DLS lesion group animals across FR5-FR7 training | Repeated measures mixed effects model (compound symmetry, with Greenhouse-Geisser correction) | Chunk duration [peak probability density (seconds)] | sequence #<br>[chronological order of sequence]<br>- *within-subjects*<br><br>group<br>[Sham or DLS lesion]<br>- *between subjects* | *simple effect* (sequence #)<br>FR5-7: $F_{(8.322, 97.46)} = 2.118$<br><br>*main effect* (group)<br>FR5-7: $F_{(1,14)} = 3.048$<br><br>*interaction*<br>(sequence #*group)<br>FR5-7: $F_{(159, 1862)} = 1.465$ | $p < 0.05$<br><br>$p > 0.05$<br><br>$p < 0.001$ |

| Figure | Analysis | Test | Dependent Variable/s | Independent Variable/s | Statistics | P value |
|---|---|---|---|---|---|---|
| **F3-FS1I** | Duration of whole successful sequences as they occur chronologically in Sham and DLS lesion group animals across FR3-FR5-FR7 training | Repeated measures mixed effects model (compound symmetry, with Greenhouse-Geisser correction) | Chunk duration [peak probability density (seconds)] | sequence # [chronological order of sequence] - *within-subjects*<br><br>group [Sham or DLS lesion] - *between subjects* | *simple effect* (sequence #)<br>FR3-7: $F_{(3.260, 42.25)} = 1.304$<br><br>*main effect* (group)<br>FR3-7: $F_{(1,14)} = 1.946$<br><br>*interaction* (sequence #*group)<br>FR3-7: $F_{(239, 3097)} = 1.192$ | $p > 0.05$<br><br>$p > 0.05$<br><br>$p < 0.001$ |
